# Antibodies to the SARS-CoV-2 receptor-binding domain that maximize breadth and resistance to viral escape

**DOI:** 10.1101/2021.04.06.438709

**Authors:** Tyler N. Starr, Nadine Czudnochowski, Fabrizia Zatta, Young-Jun Park, Zhuoming Liu, Amin Addetia, Dora Pinto, Martina Beltramello, Patrick Hernandez, Allison J. Greaney, Roberta Marzi, William G. Glass, Ivy Zhang, Adam S. Dingens, John E. Bowen, Jason A. Wojcechowskyj, Anna De Marco, Laura E. Rosen, Jiayi Zhou, Martin Montiel-Ruiz, Hannah Kaiser, Heather Tucker, Michael P. Housley, Julia di Iulio, Gloria Lombardo, Maria Agostini, Nicole Sprugasci, Katja Culap, Stefano Jaconi, Marcel Meury, Exequiel Dellota, Elisabetta Cameroni, Tristan I. Croll, Jay C. Nix, Colin Havenar-Daughton, Amalio Telenti, Florian A. Lempp, Matteo S. Pizzuto, John D. Chodera, Christy M. Hebner, Sean P.J. Whelan, Herbert W. Virgin, David Veesler, Davide Corti, Jesse D. Bloom, Gyorgy Snell

**Affiliations:** Basic Sciences Division, Fred Hutchinson Cancer Research Center, Seattle, WA 98109, USA; Vir Biotechnology, San Francisco, CA 94158, USA; Humabs BioMed SA, a subsidiary of Vir Biotechnology, 6500 Bellinzona, Switzerland; Department of Biochemistry, University of Washington, Seattle, WA 98195, USA; Department of Molecular Microbiology, Washington University School of Medicine, St. Louis, MO 63110, USA; Department of Genome Sciences, University of Washington, Seattle, WA 98195, USA; Computational and Systems Biology Program, Sloan Kettering Institute, Memorial Sloan Kettering Cancer Center, New York, NY 10065, USA; Tri-Institutional PhD Program in Computational Biology and Medicine, Weill Cornell Graduate School of Medical Sciences, New York, NY 10065, USA; Cambridge Institute for Medical Research, Department of Haematology, University of Cambridge, Cambridge, CB2 0XY, UK; Molecular Biology Consortium, Advanced Light Source, Lawrence Berkeley National Laboratory, Berkeley, CA 94720, USA; Department of Pathology and Immunology, Washington University School of Medicine, Saint Louis, MO 63110, USA; Department of Internal Medicine, UT Southwestern Medical Center, Dallas, TX 75390, USA; Howard Hughes Medical Institute, Seattle, WA 98109, USA

## Abstract

An ideal anti-SARS-CoV-2 antibody would resist viral escape^1–3^, have activity against diverse SARS-related coronaviruses^4–7^, and be highly protective through viral neutralization^8–11^ and effector functions^12,13^. Understanding how these properties relate to each other and vary across epitopes would aid development of antibody therapeutics and guide vaccine design. Here, we comprehensively characterize escape, breadth, and potency across a panel of SARS-CoV-2 antibodies targeting the receptor-binding domain (RBD), including S309^4^, the parental antibody of the late-stage clinical antibody VIR-7831. We observe a tradeoff between SARS-CoV-2 *in vitro* neutralization potency and breadth of binding across SARS-related coronaviruses. Nevertheless, we identify several neutralizing antibodies with exceptional breadth and resistance to escape, including a new antibody (S2H97) that binds with high affinity to all SARS-related coronavirus clades via a unique RBD epitope centered on residue E516. S2H97 and other escape-resistant antibodies have high binding affinity and target functionally constrained RBD residues. We find that antibodies targeting the ACE2 receptor binding motif (RBM) typically have poor breadth and are readily escaped by mutations despite high neutralization potency, but we identify one potent RBM antibody (S2E12) with breadth across sarbecoviruses closely related to SARS-CoV-2 and with a high barrier to viral escape. These data highlight functional diversity among antibodies targeting the RBD and identify epitopes and features to prioritize for antibody and vaccine development against the current and potential future pandemics.

## Main Text

The most potently neutralizing antibodies to SARS-CoV-2—including those in current clinical use^14–16^ and dominant in polyclonal sera^17,18^—target the spike receptor-binding domain (RBD). Mutations in the RBD that reduce binding by therapeutic and vaccine-elicited antibodies have recently emerged across multiple SARS-CoV-2 lineages^19–23^, highlighting the need for antibodies and vaccines that are robust to viral escape. Longer term, identification of antibodies with broad activity across SARS-related coronaviruses (sarbecoviruses) would be useful to combat potential future sarbecovirus outbreaks^6^. We have previously described an antibody, S309^4^, that exhibits potent effector functions and neutralizes all current SARS-CoV-2 isolates^24^ and the divergent sarbecovirus SARS-CoV-1, and forms the basis for an antibody therapy (VIR-7831) that has recently demonstrated clinical efficacy for treatment of COVID-19^25^. The continued development of antibodies that are resilient to SARS-CoV-2 and sarbecovirus evolution would be aided by a systematic understanding of the relationships among RBD epitope, SARS-CoV-2 neutralization potency, resistance to viral escape, and breadth of sarbecovirus cross-reactivity. Here we address this question by comprehensively characterizing a diverse panel of antibodies including S309 using deep mutational scanning, pan-sarbecovirus binding assays, *in vitro* selection of viral escape, and biochemical and structural analyses.

### Potency, escapability, and breadth in a diverse panel of RBD antibodies

We identified a panel of anti-SARS-CoV-2 antibodies with distinct properties (**Fig. 1a**, **Extended Data Table 1**), including six antibodies newly described in this study. These antibodies bind different RBD epitopes, including epitopes that overlap the receptor-binding motif (RBM) and epitopes in the non-RBM “core” of the RBD. The antibody panel spans a range of neutralization potencies (as measured with both authentic SARS-CoV-2 and spike-pseudotyped VSV particles) and binding affinities (**Fig. 1a, Extended Data Fig. 1a-c**).

**Fig. 1.**
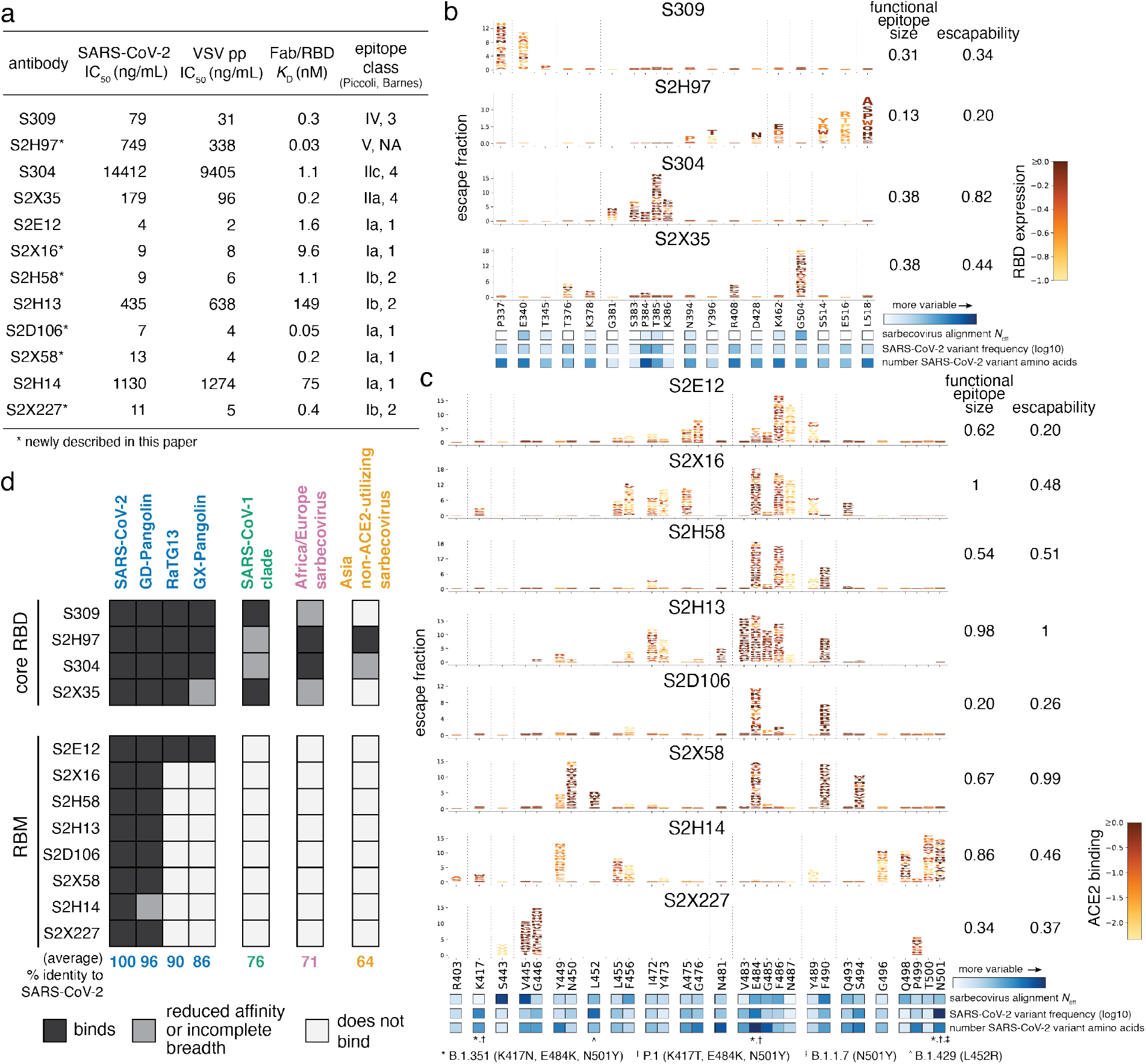
Potency, escapability, and breadth in a diverse panel of RBD antibodies. **a**, For a panel of 12 SARS-CoV-2 antibodies, summary of neutralization potency (authentic SARS-CoV-2-NLuc [*n*=3] and SARS-CoV-2 spike-pseudotyped VSV particles [*n* = 3 to 8] on Vero E6 cells, Extended Data Fig. 1a,b), 1:1 Fab:RBD binding affinities (SPR, Extended Data Fig. 1c), and epitope class according to the schemes of Piccoli et al.^17^ and Barnes et al.^9^ See Extended Data Table 1 for additional antibody details. Binding affinities for previously described antibodies measured in Piccoli et al. (S304, S2X35, S2H13, S2H14)^17^, Tortorici et al. (S2E12)^8^ and Cathcart et al. (S309)^24^ **b,c**, Complete maps of mutations that escape binding by antibodies targeting the core RBD (**b**) or receptor-binding motif [RBM] (**c**), as determined by a yeast-display deep mutational scanning method (Extended Data Fig. 2). In each map, the height of a letter indicates that mutation’s strength of escape from antibody binding. Letters are colored by their effects on folded RBD expression (**b**) or ACE2 binding affinity (**c**) [scale bars, right], as determined in our prior deep mutational scans^28^. See Extended Data Fig. 3 for escape maps colored by the opposite functional property. For each antibody, the relative “functional epitope size” and “escapability” are tabulated at right (see Methods for details). Heatmaps at bottom illustrate variability of each position within the sarbecovirus alignment or among globally sampled SARS-CoV-2 mutants. See Extended Data Fig. 4a for mutation-level variability and escape among observed SARS-CoV-2 mutants. Interactive escape maps and structural visualizations can be found at: https://jbloomlab.github.io/SARS-CoV-2-RBD_MAP_Vir_mAbs. **d**, Breadth of sarbecovirus binding by each antibody, summarizing comprehensive pan-sarbecovirus RBD yeast-display, ELISA, mammalian-display, and SPR binding assays. See Extended Data Fig. 5 for all data and phylogenetic definition of RBD clades. Black indicates an antibody binds an RBD or all RBDs within a clade with binding strength similar to SARS-CoV-2, gray indicates binding is reduced in affinity, or not all homologs within a clade are strongly bound, and white indicates no binding detected to a homolog or within a clade. The percent identity between RBD amino acid sequences with SARS-CoV-2 (or average % identity for clades) is shown below each column.

We used a deep mutational scanning platform leveraging yeast-surface displayed RBD mutant libraries to comprehensively map how all single amino-acid mutations in the SARS-CoV-2 RBD affect binding by each antibody^3^ (**Fig. 1b,c** and **Extended Data Figs. 2, 3**). The resulting maps reveal variation in the sites of escape across antibodies. Some antibodies have narrowly focused functional epitopes (a “functional epitope” comprises a set of residues where mutations abolish binding^26,27^), with binding-escape mutations at just a few key residues (e.g., S309, S2D106), while other antibodies have wider functional epitopes with binding-escape mutations at many sites (e.g., S2H13; see tabulations of “functional epitope size” to right of the logoplots in Fig. 1b,c). We previously measured how all RBD mutations affect ACE2 binding affinity and expression of folded RBD^28^, which we project onto our escape maps (letter colors in Fig. 1b,c, **Extended Data Fig. 3**). We used the combined measures of how mutations affect antibody binding and RBD function to compute the “escapability” of each antibody, which is a measure of the extent to which mutations that escape antibody binding are functionally tolerated (see tabulations to the right of the logoplots in Fig. 1b,c; discussed more below). We also investigated the sensitivity of each antibody to mutations among SARS-CoV-2 sequences reported in the GISAID database as of 4 March, 2021 (heatmap below logoplots in Fig. 1b,c; **Extended Data Fig. 4a**), and found that some antibodies are much more affected by natural SARS-CoV-2 mutations than others, including mutations found in rapidly spreading SARS-CoV-2 variants of concern^29–32^.

We next extended our deep mutational scanning platform to measure binding of each antibody to all 45 known sarbecovirus RBDs (**Fig. 1d** and **Extended Data Fig. 5**). The four antibodies that bind the core RBD exhibit cross-reactive binding to the RBDs from SARS-CoV-1 and related ACE2-utilizing bat sarbecoviruses (average 76% amino acid identity to SARS-CoV-2) and from sarbecoviruses in Europe and Africa (average 71% amino acid identity). Antibodies S304 and S2H97 also bind the most divergent clade of non-ACE2-utilizing RBDs from Asia that have an average of only 64% amino acid identity with SARS-CoV-2. S2H97 exhibits notably tight binding even to these most divergent RBDs (**Extended Data Fig. 5a,f**), making S2H97 the broadest pan-sarbecovirus RBD antibody described to date. Consistent with these binding results, S2H97 can neutralize VSV pseudotyped with diverse SARS-CoV-2- and SARS-CoV-1-related spikes (**Extended Data Fig. 5g**). In contrast, antibodies that bind epitopes within the RBM exhibit more limited cross-reactivity, typically binding only SARS-CoV-2 and the closely related GD-Pangolin-CoV RBD. This is consistent with the high sequence diversity within the RBM, which tolerates a broad range of amino acid substitutions while retaining the capacity to bind ACE2 (see **Fig. 4a**)^22,28^. However, unique among the RBM antibodies we evaluated, S2E12 binds all RBDs within the SARS-CoV-2 clade extending out to the GX-Pangolin-CoV RBD, which shares 86% amino acid identity with SARS-CoV-2. This shows that even within the evolutionarily plastic RBM there are epitopes that enable greater breadth than others.

### Relationships between antibody epitope, neutralization potency, escapability, and breadth

We next examined how antibody neutralization potency, mutant escapability, and sarbecovirus breadth relate to one another and to RBD epitope. We used our experimentally determined escape maps (**Fig. 1b,c**), together with comparable published maps for other RBD antibodies^3,21,23,33^, to project the antibodies into a two-dimensional space based on similarities in sites of binding-escape mutations (**Fig. 2a**). Although this projection is generated solely from escape maps, it recapitulates the structural space of antibody epitopes (see structures arrayed around Fig. 2a,b). In contrast to the coarse classification of epitope bins, we observe a smooth continuum of antibody epitopes that tile across the RBM (see arc from LY-CoV016 through S2X58 to REGN10987). The projection also identifies antibodies binding to unique surfaces on the RBD, such as S2H97, which binds the lateral surface of the core RBD below the receptor-binding ridge, a surface that packs with the spike N-terminal domain (NTD) of a neighboring protomer in the closed spike trimer.

**Fig. 2.**
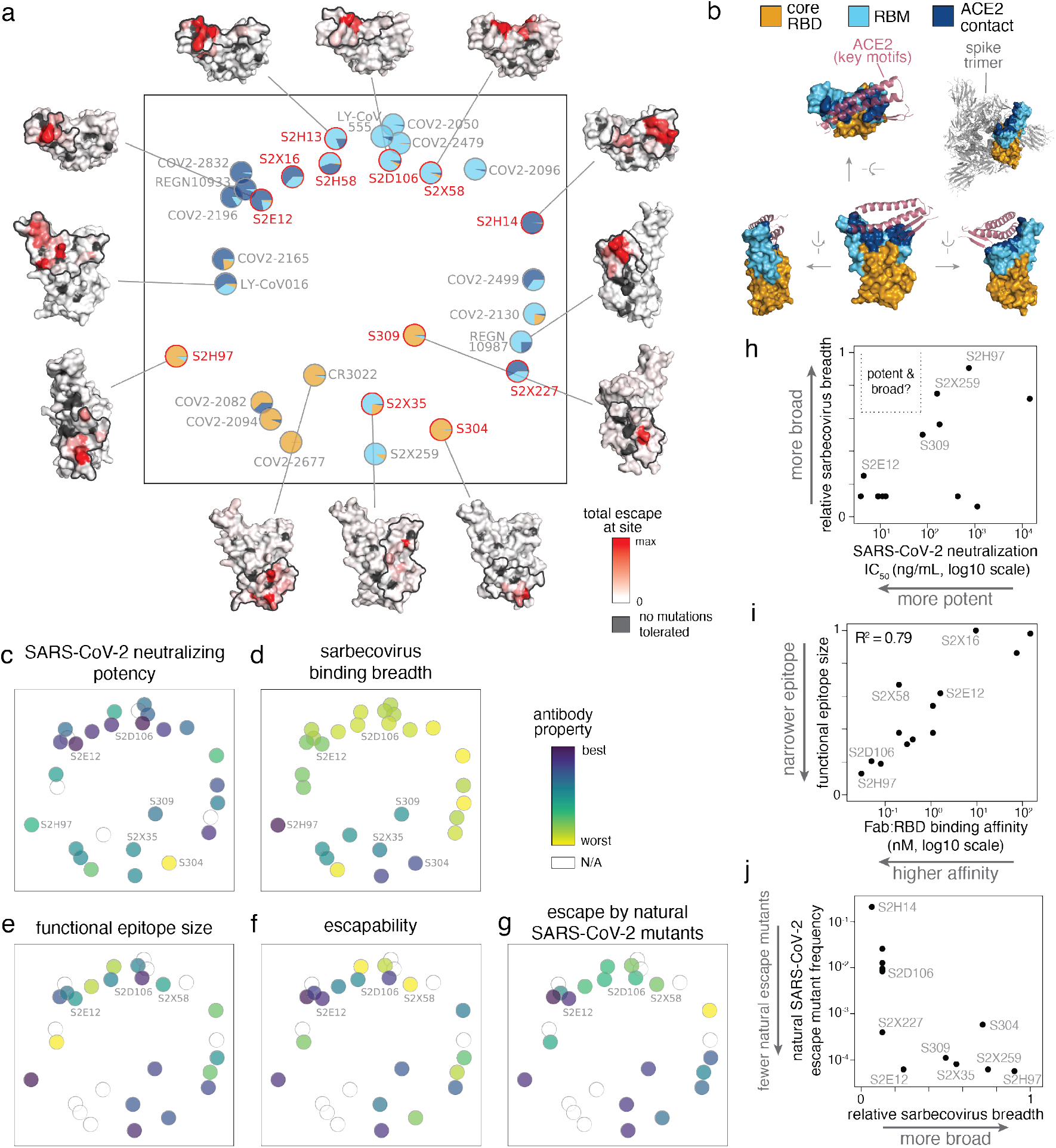
Relationship between antibody epitope, potency, breadth, and escapability. **a**, Multidimensional scaling projection of antibody epitopes based on similarities in sites of binding-escape as mapped in this (red) or prior (gray) studies. Pie charts illustrate the RBD sub-domains where mutations confer escape for each antibody [see key, (**b**)]. Structural projections of escape from representative antibodies are arrayed around the perimeter (scale bar, bottom right), with gray outlines tracing the structural footprint for antibodies with solved structures. **b**, Structural key for (**a**), illustrating the relative angles of structural views, the classification of sub-domains, and the context of ACE2 binding (only the interacting structural elements are shown) and spike quaternary structure. **c-g**, Projected epitope space from (**a**) annotated by antibody properties. For each property, antibodies are colored such that purple (scale bar, upper-right) reflects the most desirable antibody: most potent neutralization (IC50, log10 scale), highest breadth, narrowest functional epitope size, lowest escapability, and lowest natural frequency of escape mutants (log10 scale). See Methods for definition of each property, and Extended Data Fig. 6a for quantitative scales of each property and all of their pairwise correlations. **h**, Relationship between sarbecovirus breadth and SARS-CoV-2 neutralization potency for antibodies characterized in this study and S2X259 (see accompanying paper, Tortorici et al.). **i**, Relationship between functional epitope size and RBD binding affinity for antibodies characterized in this study and S2X259. Note that binding-escape selections were calibrated independently for each antibody (Extended Data Fig. 2, Methods), so this correlation is not a simple consequence of high-affinity antibodies having fewer mutations that reduce affinity below some global threshold of escape applied universally to all antibodies. **j**, Relationship between natural SARS-CoV-2 mutation escape (summed frequency of all binding-escape mutations on GISAID, Extended Data Fig. 4a) and sarbecovirus breadth for antibodies characterized in this study and S2X259.

We annotated our projection of epitope space by antibody properties such as *in vitro* neutralization potency, breadth, and escapability (**Fig. 2c-g**; see **Extended Data Fig. 6a** for all correlations). The most potently neutralizing antibodies (e.g., S2E12, S2D106) bind epitopes in the RBM, while antibodies targeting the core RBD are less potently neutralizing (**Fig. 2c**). It is important to note that RBD antibodies can protect *in vivo* through other mechanisms beyond neutralization^12,13,24^. Antibodies with broad sarbecovirus binding universally target the core RBD (**Fig. 2d**), illustrating a general tradeoff between breadth of sarbecovirus binding (i.e., conservation within an epitope) and potency of SARS-CoV-2 neutralization (Fig. 2h). Nonetheless, some cross-reactive antibodies exhibit moderate *in vitro* neutralization potency (e.g., S309, S2X259 [see accompanying paper Tortorici et al.]), and the highly potent RBM-directed antibody S2E12 exhibits moderate breadth, highlighting the existence of antibodies with simultaneous neutralization potency, breadth, and resistance to mutational escape.

In contrast to the relationships between breadth and potency, the “size” of an antibody’s functional epitope (the sum of mutation escape fractions, **Fig. 1b,c**) is not strongly influenced by the epitope’s structural location (**Fig. 2e**)—instead, narrower functional epitopes are associated with higher Fab:RBD binding affinity (**Fig. 2i**). However, the relationship between an antibody’s functional epitope size and its “escapability,” which integrates how escape mutations affect RBD folding and ACE2 affinity, is influenced by variation in these functional constraints across the RBD structure. For example, S2E12 has a functional epitope that is similarly wide as that of S2X58, while S2D106 exhibits a narrower functional epitope (**Figs. 1c, 2e**). However, S2E12 focuses on functionally constrained ACE2 contact residues while S2D106 and S2X58 target functionally tolerant sites that vary across SARS-CoV-2 sequences (**Fig. 1c**). Therefore, despite its wider functional epitope, S2E12 exhibits low escapability (**Fig. 2f**) and is affected by fewer observed SARS-CoV-2 mutations than S2D106 (**Fig. 2g** and **Extended Data Fig. 4a**)^19^. The low escapability of S2E12 is mirrored in its unique breadth among RBM-directed antibodies: S2E12 exhibits binding across sarbecoviruses closely related to SARS-CoV-2, including RaTG13 and GX-Pangolin-CoV (**Fig. 1d**). The broadest antibodies that bind the core RBD also have low escapability (**Fig. 2f,g,j**), because they bind the RBD with high affinity (**Extended Data Fig. 6a**) at sites that are constrained with respect to RBD folding (**Fig. 1b**). Taken together, our results suggest that screening antibodies for high-affinity binding and sarbecovirus breadth—even as modestly as to GX-Pangolin-CoV—identifies antibodies that are robust to ongoing SARS-CoV-2 evolution (**Fig. 2j**), whereas screening for neutralization potency alone will uncover RBM-directed antibodies independent of their escapability.

To further support our inferences of escapability, we performed viral-escape selection experiments to identify spike-expressing VSV mutants that emerge under selection with 7 of the antibodies from the panel (**Fig. 3a** and **Extended Data Fig. 6b,c**). These selections validate the sites of escape identified in deep mutational scanning profiles (**Fig. 1b,c**) and the quantitative measure of antibody escapability derived from the deep mutational scans (**Fig. 3b**). For example, S2X58 selects escape viruses carrying mutations at many sites, while S2D106 escape is associated with mutations at the two sites identified by deep mutational scanning (**Fig. 1c**). S2E12 selects viral mutants at just a handful of sites, and these mutations reduce ACE2 binding affinity (**Fig. 3a**). Escape mutations selected by S2X58 and S2D106 (e.g. L452R and E484K) are present in SARS-CoV-2 variants of concern^30–32^, but S2E12 escape mutations are not. This fact could reflect both higher functional constraint on the S2E12 functional epitope^28^ and a relative rarity of S2E12-like antibodies in polyclonal sera leading to little antigenic pressure at these sites.

**Fig. 3.**
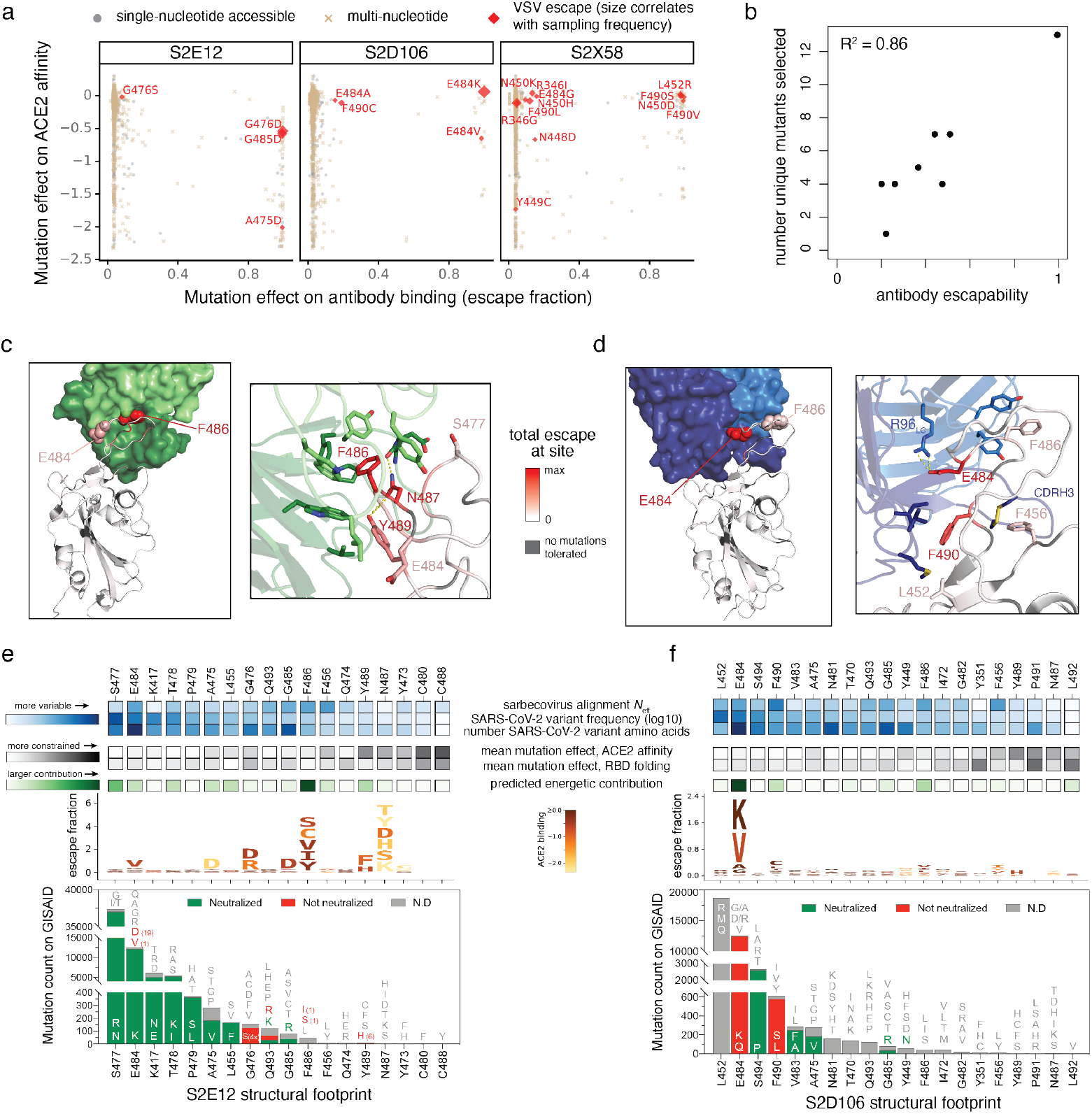
Structural basis for variation in antibody escapability. **a**, Escape mutations identified in spike-expressing VSV passaged in the presence of monoclonal antibody, illustrated with respect to their effects on antibody (x-axis) and ACE2 (y-axis) binding. Amino acid mutations are coded by whether they are accessible via single-nucleotide mutation from the wildtype spike gene sequence used in the VSV selections (Wuhan-Hu-1+D614G). See additional selections, Extended Data Fig. 6b. **b**, Correlation between the number of unique mutations selected across viral escape selection experiments and antibody escapability as tabulated in Fig. 1b,c, plus S2X259 (see accompanying paper, Tortorici et al.). **c,d**, Structures of S2E12 (**c**) and S2D106 (**d**) bound to RBD. RBD sites are colored by escape (scale bar, center). Antibody heavy chains colored dark green (S2E12) or blue (S2D106), with light chains in lighter shades. Righthand images show zoomed-in context of key structural features impacting antibody escapability. **e**,**f**, Integrative genetic and functional features of the structural epitopes of S2E12 (**e**) and S2D106 (**f**). Sites within the structural footprint of each antibody (5 Å cutoff) are ordered by the frequency of observed mutants among SARS-CoV-2 sequences present on GISAID. Heatmaps illustrate evolutionary variability (blue), functional constraint from prior deep mutational scanning measurements (gray) and predicted energetic contribution of a residue derived from the structures (green). Logoplots illustrate escape from antibody binding as in Fig. 1c, but only showing amino acid mutations accessible via single-nucleotide mutation from Wuhan-Hu-1 for comparison with (**a**). Barplots illustrate the frequency of SARS-CoV-2 mutants at each position and their validated effects on antibody neutralization measured in spike-pseudotyped VSV particles with Vero E6 cells (red, >3-fold increase in IC50 due to mutation).

### Structural basis for antibody robustness to escape

To understand the structural basis of escape for mAbs targeting the RBM, we focused on S2E12 and S2D106, which have overlapping epitopes but different escape profiles. We determined structures of S2E12 (X-ray crystallography, 2.93 Å resolution) and S2D106 (cryoEM, 4.0 Å resolution local refinement of the RBD/Fv region) bound to RBD (**Fig. 3c,d, Extended Data Fig. 7a-d and Extended Data Table 2**). We also integrated evolutionary, functional, and structural details for the sites in each antibody’s structural footprint (**Fig. 3e,f**). S2E12 and S2D106 bind the receptor-binding ridge, with 8 contact residues shared between the footprints of S2E12 (18 contacts, 5Å distance cutoff) and S2D106 (20 contacts). S2E12 binding is oriented toward extensive packing of the ACE2-contact residue F486_RBD_ within a buried cavity lined by aromatic residues at the antibody light/heavy-chain interface. The functionally constrained residue N487_RBD_ also nestles in this pocket, where it forms polar contacts with the antibody backbone (**Fig. 3c**). Sites within the S2E12 structural footprint that exhibit less functional constraint and more variation among SARS-CoV-2 isolates (e.g. E484, S477) are located at the periphery of the interface, explaining the robustness of S2E12 binding to SARS-CoV-2 mutations with high natural variant frequencies^19^ (**Fig. 3e**). This structural interface also explains the breadth of S2E12 binding to RaTG13 and GX-Pangolin-CoV (**Fig. 1d**), as the F486L mutation present in these RBDs retains the central hydrophobic packing at the interface between the S2E12 Fab and the RBD (**Fig. 2c**).

In contrast to S2E12, S2D106 binding is anchored by a buried polar contact between E484_RBD_ and R96_LC_, in addition to nonpolar contacts between F490_RBD_ and residues in the heavy chain CDR2 loop (**Fig. 3d**). Although the long heavy chain CDR3 packs intimately across the surface of the RBD through shape complementarity, including with constrained residues such as F456_RBD_, there are no crucial CDRH3:RBD contacts that are sensitive to mutation. S2D106 escape is therefore highly focused on E484 and F490, which are functionally tolerant and exhibit substantial variation among SARS-CoV-2 sequences (**Fig. 2e**), including emerging variants of concern. This comparison between S2E12 and S2D106 highlights how a subtle change in the focus of the RBD:antibody interaction impacts the robustness of each antibody to viral escape, including among natural SARS-CoV-2 mutants.

### Structural basis for antibody breadth

To understand the structural basis for breadth of sarbecovirus binding, we determined the structures of S2H97 (X-ray crystallography, 2.65 Å resolution) and S2X35 (X-ray crystallography, 1.83 Å resolution) bound to RBD (**Fig. 4a** and **Extended Data Table 2**) and a cryoEM structure of S2H97 bound to SARS-CoV-2 S (**Extended Data Fig. 7e-h**), and analyzed previously published structures of S309 and S304 bound to RBD^4,17^. This panel of broad antibodies emphasizes the core RBD as a general target of broad antibody binding due to its conservation throughout sarbecovirus evolution, reflected in the diverse core RBD surfaces targeted by these antibodies (**Fig. 4a**).

**Fig. 4.**
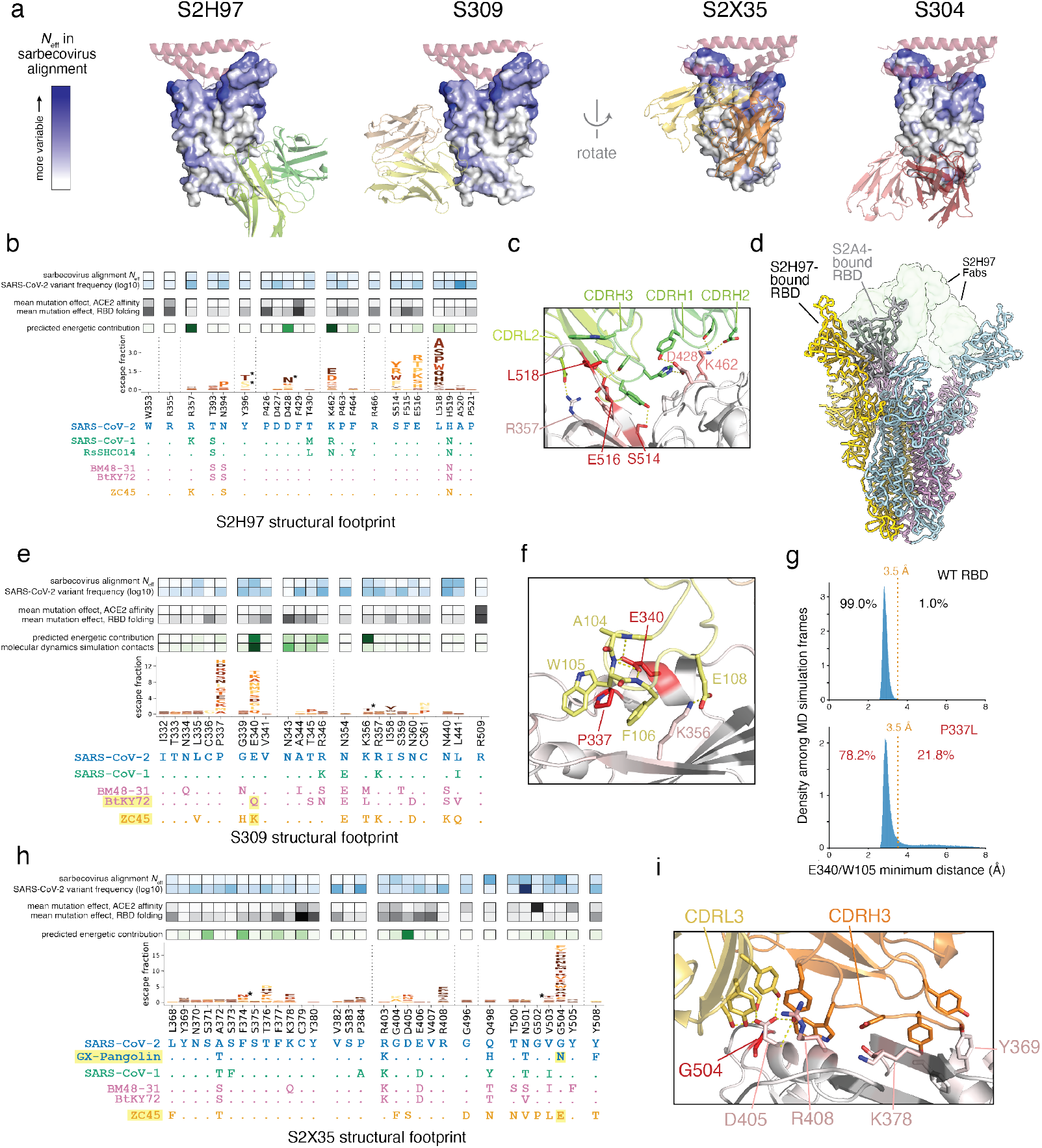
Structural basis for broad sarbecovirus binding. **a**, Overview of the surfaces targeted by broadly binding RBD antibodies. RBD surface is colored by site variability in the sarbecovirus alignment (effective number of amino acids, scale bar at left). ACE2 (key motifs) shown in transparent cartoon. Antibody variable domains shown as cartoon, with darker shade indicating the heavy chain. **b**, Integrative genetic and structural features of the S2H97 structural epitope (5 Å cutoff). Heatmap details and scale bars as in Fig. 3e,f. Logoplots are colored by mutation effects on folded RBD expression (see scale bar, Fig. 1b). Asterisks in logoplot indicate escape mutations that introduce N-linked glycosylation motifs (NxS/T). Below the logoplot is a selection of aligned sarbecovirus RBDs (sequenced colored by clade as in Fig. 1d, Extended Data Fig. 5a). **c**, Zoomed in view of the S2H97/RBD interface, with important contact and escape residues labeled. **d**, CryoEM structure of S2H97-bound SARS-CoV-2 S. Spike protomers are shown in yellow, blue, and pink, and S2H97 Fabs in transparent green surface. S2A4-bound spike protomer from PDB 7JVC^17^ is shown in gray and aligned to the yellow subunit, indicating the additional extent of RBD opening necessary to access the S2H97 epitope compared to a class II antibody. **e**, Integrative features of the S309 structural epitope, details as in (**b**). An additional row in the heatmap overlay reflects the proportion of all close S309/RBD contacts (<3.5 Å) made by each residue during molecular dynamics simulation. Highlighted sarbecoviruses identify those that escape S309 binding, and highlighted mutation in the alignment is the likely contributor according to our escape map. **f**, Zoomed in view of the S309/RBD interface, with important contact and escape residues labeled. **g**, Molecular dynamics simulations of the S309/RBD structure. Histograms show the distribution of minimum distance between E340RBD and W105HC heavy atoms across 1-ns frames during the simulation of the unmutated (top, 42-μs simulation) and P337L mutated (bottom, 91-μs simulation) RBD bound to S309. Orange line reflects the 3.5 Å distance cutoff used to define close contact. Percentage of frames in which E340 and W105 are or are not in close contact is labeled. See Extended Data Fig. 8c for the occupancy of other S309:RBD contacts across the simulations. **h**, Integrative features of the S2X35 structural epitope, details as in (b). **i**, Zoomed in view of the S2X35/RBD interface, with important contact and escape residues labeled.

The exceptional cross-reactivity of S2H97 is achieved by targeting a previously undescribed antigenic site on the RBD surface that is highly conserved across sarbecoviruses, which we designated site V (**Fig. 4a,b**). S2H97 binding is facilitated by close packing of the heavy chain CDR3 into an RBD crevice at the center of the epitope, in addition to key polar contacts with all three heavy chain CDR loops and the light chain CDR2 (**Fig. 4c**). The RBD surface bound by S2H97 is functionally constrained (Fig. 4b), reflected in the deleterious effects of mutations within this surface for folded RBD expression^28^, and this constraint is likely enhanced by quaternary packing of this surface with the NTD in the closed spike trimer (**Extended Data Fig. 8a**). S2H97 binding requires extensive opening of the RBD to unmask this epitope (**Fig. 4d**), even more so than for antibodies targeting antigenic site II, which may explain an apparent rarity of antibodies targeting this surface and the relatively low neutralization potency of S2H97 despite its high binding affinity to the isolated RBD.

S309 binding exhibits focused engagement of a small number of residues, thereby tolerating low-level sequence diversity within its structural footprint. S309 escape is mediated by mutations at RBD residues E340 and P337 which are respectively engaged in electrostatic and van der Waals interactions with CDRH3 (**Fig. 4e,f**, and **Extended Data Fig. 8b,c**). Molecular dynamics simulations (42 μs) further highlight E340 as a key antibody-contact residue, forming persistent contacts with three consecutive CDRH3 backbone amides (**Fig. 4e,f** and **Extended Data Fig. 8c**). This is in contrast to e.g. K356_RBD_, which forms polar contacts to S309 that are more transient during the simulation (**Fig. 4e** and **Extended Data Fig. 8c**). Although P337_RBD_ does not form a critical antibody contact, mutation of this helix-start proline appears to escape S309 binding either through steric hindrance with the W105 and F106 side chains or indirectly via destabilization of E340_RBD_ (**Fig. 4g** and **Extended Data Fig. 8c**). Consistent with this focused binding interaction, S309 can neutralize circulating SARS-CoV-2 mutants^24^ (**Extended Data Fig. 8b**) and bind divergent RBDs such as BM48-31 (**Extended Data Fig. 5f**), which differs from SARS-CoV-2 and SARS-CoV-1 at 8 residues within the S309 footprint (including e.g. K356M) but retains E340.

S2X35 binds a large interface through long light- and heavy-chain CDR3 loops (**Fig. 4h,i**). The S2X35 footprint extends from packing of aromatic residues within the ACE2-distal core RBD (e.g., Y369) to close packing of the S2X35 CDRH3, CDRL3, and CDRL1 with RBD residues D405, R408, and the G504 loop near the RBM. In contrast to e.g., S309 or S2E12 where the primary sites of escape alter crucial side chains for antibody binding, the primary site of S2X35 escape is G504_RBD_, where mutations instead mediate escape due to introduction of steric clash with D405_RBD_ and the S2X35 light chain—not because of any key role of G504 itself. The residue with the strongest interaction from the crystal structure is D405_RBD_, but escape at this position occurs only via biochemically dramatic amino acid changes (e.g., D405K). The restricted escape profile for S2X35 suggests less energetic dependence on any single RBD side chain for its binding, enhancing its breadth and robustness to escape. Taken together, our structural survey illustrates diverse molecular mechanisms of RBD engagement that are compatible with sarbecovirus breadth.

### Principles for optimizing antibody and vaccine development

Ongoing SARS-CoV-2 evolution^22,29–32^, long-term antigenic evolution of other human coronaviruses^34,35^, and the spillover potential of diverse sarbecovirus lineages^6,7^ indicate the importance of developing antibodies and vaccines that are robust to viral evolution. In this work, we identify antibody and epitope features which can guide this process. Although *in vitro* neutralization potency is often prioritized for lead selection, our results suggest this will bias antibodies toward RBM epitopes, many of which are poorly conserved in the short-term evolution of SARS-CoV-2^22^ and the long-term evolution of sarbecoviruses^7^. Our results suggest that additional prioritization of high affinity binding and at least a moderate degree of sarbecovirus breadth will yield antibodies with improved resistance to viral evolution. For example, the high-affinity, pan-sarbecovirus antibody S2H97 is extremely robust to global SARS-CoV-2 sequence variation. The highly potent RBM-directed antibody S2E12 binds the modestly diverged GX-Pangolin-CoV RBD, and correspondingly exhibits lower vulnerability to circulating mutations than many other RBM antibodies^19^. Therefore, our comprehensive mapping highlights the existence of antibodies targeting multiple epitopes that are broad and resistant to viral escape.

The global emergence of variants of concern (VOC) has been a defining feature in the progression of the current pandemic^29–32^. Many common mutations in VOCs occur in the RBM (e.g., residues E484, K417 and L452) and impact binding by polyclonal serum and some therapeutic antibodies^19–23^. Five of the eight RBM-directed antibodies in our panel show vulnerability to mutations at position 484, which is also a concern for several late-stage antibodies^19–21^. Comprehensive maps of escape enable identification of antibodies robust to current SARS-CoV-2 variation and can guide the targeted use of approved antibody therapeutics in response to local viral variants. We cannot predict exactly which mutations will next rise to prominence as SARS-CoV-2 continues to evolve, but it seems likely that they will include additional RBM mutations that strongly reduce recognition by infection- and vaccine-elicited antibody immunity^1,2,17,18,22^. Therefore, antibody and epitope discovery efforts focused on breadth^4,5^, aided by high-resolution differentiation among antibody epitopes as generated herein, can inform the development of countermeasures with greater robustness to immune escape in the current SARS-CoV-2 pandemic and utility for potential future sarbecovirus spillovers.

## Acknowledgements

We thank Isaac Hoffman for assistance in refinement of crystal structures, Gregory R. Bowman, Joseph Coffland, and Peter K. Eastman for developing and maintaining the Folding@home infrastructure, Amazon Web Services for Folding@home infrastructure support, the Folding@home volunteers who contributed their computational resources to this project (FAH Project 17336-17340), Rafal P. Wiewiora and Sukrit Singh for their assistance with Folding@home, and Aoife M. Harbison and Elisa Fadda for assistance in glycan modeling. This work was supported by the Fred Hutch Flow Cytometry and Genomics facilities, the Fred Hutch Scientific Computing group supported by ORIP grant S10OD028685, and the University of Washington Arnold and Mabel Beckman cryoEM center. This work was supported by the NIH/NIAID (R01AI127893 and R01AI141707 to JDB, DP1AI158186 and HHSN272201700059C to DV, T32AI083203 to AJG), the NIH/NIGMS (R01GM120553 to DV, and grant R01GM121505 and R01GM132386 to JDC), the NIH/NCI (P30CA008748 to JDC), the National Science Foundation (NSF CHI-1904822 to JDC), the Damon Runyon Cancer Research Foundation (TNS), the Gates Foundation (INV-004949 to JDB), a Pew Biomedical Scholars Award (DV), Investigators in the Pathogenesis of Infectious Disease Awards from the Burroughs Wellcome Fund (JDB and DV), the Wellcome Trust (209407/Z/17/Z to TIC), Fast Grants (DV), Bayer (WGG) and the Molecular Sciences Software Institute (IZ). JDB is an Investigator of the Howard Hughes Medical Institute. The Molecular Biology Consortium beamline 4.2.2 of the Advanced Light Source, a DOE Office of Science User Facility under Contract No. DE-AC02-05CH11231, is supported in part by the ALS-ENABLE program funded by the NIH/NIGMS (P30GM124169-01). Use of the Stanford Synchrotron Radiation Lightsource, SLAC National Accelerator Laboratory, is supported by the U.S. Department of Energy, Office of Science, Office of Basic Energy Sciences under Contract No. DE-AC02-76SF00515. The SSRL Structural Molecular Biology Program is supported by the DOE Office of Biological and Environmental Research, and by the NIH/NIGMS (P30GM133894).

## Author Contributions

Conceived research and designed study: TNS, NC, HWV, DC, JDB, GS. Antibody discovery: FZ, DP, MB, RM, ADM, EC, MSP, DC. Expression and purification of proteins: NC, PH, GL, NS, KC, SJ, MM. Antibody functional experiments: FZ, ZL, DP, MB, RM, JAW, ADM, LER, JZ, MMR, HK, HT, MPH, MA, ED, CHD, FAL, SPJW. Deep mutational scanning experiments and analysis: TNS, AA, AJG, ASD. Bioinformatics analysis: JDI, AT. Structure determination: NC, YJP, PH, JEB, TIC, JCN, DV, GS. Molecular dynamics simulation and analysis: WGG, IZ, JDC. Supervision: MSP, JDC, CMH, SPJW, DV, DC, JDB, GS. Wrote the initial draft: TNS, NC, DC, JDB, GS. Edited the final version: all authors.

## Declaration of Interests

NC, FZ, DP, MB, PH, RM, JAW, ADM, LER, JZ, MMR, HK, HT, MPH, JDI, GL, MA, NS, KC, SJ, MM, ED, EC, CHD, AT, FAL, MSP, CMH, HWV, DC and GS are employees of Vir Biotechnology and may hold shares in Vir Biotechnology. DC is currently listed as an inventor on multiple patent applications, which disclose the subject matter described in this manuscript. HWV is a founder of PierianDx and Casma Therapeutics. Neither company provided funding for this work or is performing related work. JCN, TIC, and DV are consultants for Vir Biotechnology Inc. The Veesler laboratory has received a sponsored research agreement from Vir Biotechnology Inc. JDC is a current member of the Scientific Advisory Boards of OpenEye Scientific Software, Interline Therapeutics, and Redesign Science. The Chodera laboratory receives or has received funding from the National Institutes of Health, the National Science Foundation, the Parker Institute for Cancer Immunotherapy, Relay Therapeutics, Entasis Therapeutics, Silicon Therapeutics, EMD Serono (Merck KGaA), AstraZeneca, Vir Biotechnology, XtalPi, Interline Therapeutics, and the Molecular Sciences Software Institute, the Starr Cancer Consortium, the Open Force Field Consortium, Cycle for Survival, a Louis V. Gerstner Young Investigator Award, and the Sloan Kettering Institute. A complete funding history for the Chodera lab can be found at http://choderalab.org/funding. The other authors declare no competing interests.

## Additional Information

Correspondence and requests for materials should be addressed to Davide Corti, Jesse Bloom, and Gyorgy Snell.

## Materials and Methods

### Isolation of peripheral blood mononuclear cells (PBMCs), plasma and sera

Samples from three SARS-CoV-2 recovered individuals, designated as donors S2H, S2D and S2X were obtained under study protocols approved by the local Institutional Review Board (Canton Ticino Ethics Committee, Switzerland). All donors provided written informed consent for the use of blood and blood components (such as PBMCs, sera or plasma). Blood drawn from donor S2X was obtained at day 48 (S2X16, S2X35 and S2X58 mAbs) and 75 (S2X227) after symptoms onset. Blood from donor S2H was obtained at day 17 (S2H13 and S2H14), day 45 (S2H58) and day 81 (S2H97) after symptoms onset. Blood from donor S2D was obtained at day 98 (S2D106) after symptoms onset.

PBMCs were isolated from blood draw performed using tubes pre-filled with heparin, followed by Ficoll density gradient centrifugation. PBMCs were either used fresh for SARS-CoV-2 Spike protein-specific memory B cell sorting or stored in liquid nitrogen for later use. Sera were obtained from blood collected using tubes containing clot activator, followed by centrifugation and stored at −80°C.

### B-cell isolation and recombinant mAb production

Discovery and initial characterization of six antibodies in our panel was previously reported (S309 and S304^4,17^, S2X35, S2H13 and S2H14^17^, and S2E12^8^), and six new antibodies are first described here (S2H97, S2X16, S2H58, S2D106, S2X58, S2X227). Starting from freshly isolated PBMCs or upon cells thawing, B cells were enriched by staining with CD19 PE-Cy7 (and incubation with anti-PE beads), followed by positive selection using LS columns. Enriched B cells were stained with anti-IgM, anti-IgD, anti-CD14 and anti-IgA, all PE labelled, and prefusion SARS-CoV-2 S with a biotinylated Avi-tag conjugated to Streptavidin Alexa-Fluor 647 (Life Technologies). SARS-CoV-2 S-specific IgG+ memory B cells were sorted by flow cytometry via gating for PE negative and Alexa-Fluor 647 positive cells. Cells were cultured for the screening of positive supernatants. Antibody VH and VL sequences were obtained by RT-PCR and mAbs were expressed as recombinant human Fab fragment or as IgG1 (G1m3 allotype) carrying the half-life extending M428L/N434S (LS) mutation in the Fc region. ExpiCHO-S (Thermo Fisher Scientific) cells were transiently transfected with heavy and light chain expression vectors as previously described^4^. Affinity purification was performed on ÄKTA Xpress FPLC (Cytiva) operated by UNICORN software version 5.11 (Build 407) using HiTrap Protein A columns (Cytiva) for full length human mAbs and CaptureSelect CH1-XL MiniChrom columns (Thermo Fisher Scientific) for Fab fragments, using PBS as mobile phase. Buffer exchange to the appropriate formulation buffer was performed with a HiTrap Fast desalting column (Cytiva). The final products were sterilized by filtration through 0.22 μm filters and stored at 4°C.

Epitope classes shown in Fig. 1a are defined as in Piccoli et al.^17^ Briefly, the classification of these epitope classes results from Octet binning experiments using structurally characterized antibodies, structural insights to define the recognition of open-only RBD and ability of antibodies to interfere with RBD binding to ACE2. In particular, site Ia is accessible only in the open state of RBD and largely overlaps with ACE2 footprint; site 1b is accessible in both open and closed RBD states and overlaps in part with ACE2 footprint; site IIa is in the core RBD (accessible only in the open RBD state) and antibodies binding to this site interfere with binding to ACE2, site IIc is also in the core RBD but targeted by antibodies that do not interfere with binding to ACE2; site IV is fully accessible on both open and closed RBDs and is defined by the footprint of S309 antibody.

### Neutralization of authentic SARS-CoV-2 by entry-inhibition assay

Neutralization was determined using SARS-CoV-2-Nluc, an infectious clone of SARS-CoV-2 (based on strain 2019-nCoV/USA_WA1/2020) which encodes nanoluciferase in place of the viral ORF7 and demonstrated comparable growth kinetics to wildtype virus^36^. Vero E6 cells (ATCC CRL-1586) were seeded into black-walled, clear-bottom 96-well plates at 2 × 10^4^ cells/well and cultured overnight at 37°C. The next day, 9-point 4-fold serial dilutions of mAbs were prepared in infection media (DMEM + 10% FBS). SARS-CoV-2-Nluc was diluted in infection media at a final MOI of 0.01 PFU/cell, added to the mAb dilutions and incubated for 30 minutes at 37°C. Media was removed from the Vero E6 cells, mAb-virus complexes were added and incubated at 37°C for 24 hours. Media was removed from the cells, Nano-Glo luciferase substrate (Promega) was added according to the manufacturer’s recommendations, incubated for 10 minutes at room temperature and the luciferase signal was quantified on a VICTOR Nivo plate reader (Perkin Elmer).

### SARS-CoV-2 spike pseudotyped VSV generation and neutralization assay

Replication defective VSV pseudovirus^37^ expressing SARS-CoV-2 spike protein were generated as previously described^38^ with some modifications. Plasmids encoding SARS-CoV-2 spike variants were generated by site-directed mutagenesis of the wild-type plasmid, pcDNA3.1(+)-spike-D19^39^. Lenti-X 293T cells (Takara, 632180) were seeded in 10-cm dishes at a density of 1e5 cells/cm^2^ and the following day transfected with 5 μg of spike expression plasmid with TransIT-Lenti (Mirus, 6600) according to the manufacturer’s instructions. One day post-transfection, cells were infected with VSV-luc (VSV-G) (Kerafast, EH1020-PM) for 1 h, rinsed three times with PBS, then incubated for an additional 24 h in complete media at 37°C. The cell supernatant was clarified by centrifugation, filtered (0.45 μm), aliquoted, and frozen at −80°C.

For VSV pseudovirus neutralization assays, Vero E6 cells (ATCC CRL-1586) were grown in DMEM supplemented with 10% FBS and seeded into clear bottom white 96 well plates (Costar, 3903) at a density of 2e4 cells per well. The next day, mAbs were serially diluted in pre-warmed complete media, mixed at a 1:1 ratio with pseudovirus and incubated for 1 h at 37°C in round bottom polypropylene plates. Media from cells was aspirated and 50 μL of virus-mAb complexes were added to cells and then incubated for 1 h at 37°C. An additional 100 μL of prewarmed complete media was then added on top of complexes and cells incubated for an additional 16-24 h. Conditions were tested in duplicate wells on each plate and at least six wells per plate contained uninfected, untreated cells (mock) and infected, untreated cells (‘no mAb control’). Virus-mAb-containing media was then aspirated from cells and 100 μL of a 1:4 dilution of Bio-glo (Promega, G7940) in PBS was added to cells. Plates were incubated for 10 min at room temperature and then were analyzed on the Envision plate reader (PerkinElmer). Relative light units (RLUs) for infected wells were subtracted by the average of RLU values for the mock wells (background subtraction) and then normalized to the average of background subtracted “no mAb control” RLU values within each plate. Percent neutralization was calculated by subtracting from 1 the normalized mAb infection condition. Data were analyzed and visualized with Prism (Version 8.4.3). IC_50_ and IC_80_ values were calculated from the interpolated value from the log(inhibitor) versus response – variable slope (four parameters) nonlinear regression with an upper constraint of < 100. Each neutralization experiment was conducted on three independent days, i.e., biological replicates, where each biological replicate contains a technical duplicate. IC_50_ values across biological replicates are presented as geometric mean. The loss or gain of neutralization potency across spike variants was calculated by dividing the variant IC_50_ by the parental (D614G) IC_50_ within each biological replicate.

### Sarbecovirus spike pseudotyped VSV neutralization by S2H97

Mammalian expression constructs (pcDNA3.1(+) or pTwist-CMV) encoding the spike proteins from various Ssarbecoviruses with a C-terminal deletion of 19 amino acids (D19) were synthesized for SARS-CoV-2 (Genbank: QOU99296.1), SARS-CoV-1 (Urbani, Genbank: AAP13441.1), hCoV-19/pangolin/Guangdong/1/2019 (GD-Pangolin-CoV, Genbank: QLR06867.1), Pangolin coronavirus Guanxi-2017 (GX-Pangolin-CoV, Genbank: QIA48623.1), bat SARS-like coronavirus WIV1 (WIV1,Genbank: AGZ48828.1). Lenti-X 293T (Takara, 632180) cells were seeded in 15 cm dishes such that the cells would reach 80% confluency after culturing overnight. The following day, cells were transfected using TransIT-Lenti (Mirus, 6600) according to the manufacturer’s instructions. One day post-transfection, cells were infected with VSV-G*ΔG-luciferase (Kerafast, EH1020-PM). The supernatant containing sarbecovirus pseudotyped VSV was collected 2 days post-transfection, centrifuged at 1000 × g for 5 minutes, aliquoted and frozen at −80°C.

For neutralization assays, cells supporting robust pseudovirus infection were seeded into clear bottom white-walled 96-well plates at 20,000 cells/well in 100 μL culture media. VeroE6 cells were used for VSV-SARS-CoV-2, VSV-SARS-CoV-1, and VSV-GD-Pangolin-CoV. BHK-21 cells stably expressing ACE2 were used for VSV-GX-Pangolin-CoV and VSV-WIV1. After culturing cells overnight, 1:3 serial dilutions of antibody were prepared in DMEM in triplicate. Pseudovirus was diluted in DMEM and added each antibody dilution such that the final dilution of pseudovirus was 1:20. Pseudovirus:antibody mixtures were incubated for 1 hour at 37°C. Media was removed from the cells and 50 μL of pseudovirus:antibody mixtures were added. One hour post-infection, 50 μL of culture media was added to wells containing pseudovirus:antibody mixtures and incubated overnight at 37°C. Media was then removed and 100 μL of 1:1 diluted DPBS:Bio-Glo (Promega, G7940) luciferase substrate was added to each well. The plate was shaken at 300 RPM at room temperature for 10 minutes after which RLUs were read on an EnSight (Perkin Elmer) microplate reader. Percent neutralization was determined by first subtracting the mean background (cells with luciferase substrate alone) RLU values of 6 wells per plate for all data points. Percent neutralization for each antibody concentration was calculated relative to no antibody control wells for each plate. Percent neutralization data were analyzed and graphed using Prism (GraphPad, v9.0.1). Absolute IC_50_ values were calculated by fitting a curve using a non-linear regression model (variable slope, 4 parameters) and values were interpolated from the curve at y=50. The geometric mean from at least two independent experiments was calculated using Excel (Microsoft, Version 16.45).

### Recombinant protein production

SARS-CoV-2 RBD WT for SPR binding assays (with N-terminal signal peptide and C-terminal thrombin cleavage site-TwinStrep-8xHis-tag) was expressed in Expi293F (Thermo Fisher Scientific) cells at 37°C and 8% CO_2_. Transfections were performed using the ExpiFectamine 293 Transfection Kit (Thermo Fisher Scientific). Cell culture supernatant was collected three days after transfection and supplemented with 10x PBS to a final concentration of 2.5x PBS (342.5 mM NaCl, 6.75 mM KCl and 29.75 mM phosphates). SARS-CoV-2 RBDs were purified using 1 or 5 mL HisTALON superflow cartridges (Takara Bio) and subsequently buffer exchanged into 1x HBS-N buffer (Cytiva) or PBS using Zeba Spin Desalting or HiPrep 26/10 desalting columns.

SARS-CoV-2 RBD WT for crystallization (with N-terminal signal peptide and ‘ETGT’, and C-terminal 8xHis-tag) was expressed similarly as described above. Cell culture supernatant was collected four days after transfection and supplemented with 10x PBS to a final concentration of 2.5x PBS. Protein was purified using a 5 ml HisTALON superflow cartridge followed by size exclusion chromatography on a Superdex 200 Increase 10/300 GL column equilibrated in 20 mM Tris-HCl pH 7.5, 150 mM NaCl. RBD was deglycosylated by overnight incubation with EndoH glycosidase at 4°C.

RBDs from other sarbecoviruses for SPR (with N-terminal signal peptide and C-terminal thrombin cleavage site-TwinStrep-8xHis-tag) were expressed in Expi293F cells at 37°C and 8% CO_2_. Cells were transfected using PEI MAX (Polysciences) at a DNA:PEI ratio of 1:3.75. Transfected cells were supplemented three days after transfection with 3 g/L glucose (Bioconcept) and 5 g/L soy hydrolysate (Sigma-Aldrich Chemie GmbH). Cell culture supernatant (423 mL) was collected seven days after transfection and supplemented with 47 mL 10x binding buffer (1 M Tris-HCl, 1.5 M NaCl, 20 mM EDTA, pH 8.0) and 25 mL BioLock (IBA GmbH) and incubated on ice for 30 min. Proteins were purified using a 5 mL Strep-Tactin XT Superflow high capacity cartridge (IBA GmbH) followed by buffer exchange to PBS using HiPrep 26/10 desalting columns (Cytiva).

Prefusion-stabilized SARS-CoV-2 spike protein for SPR (residues 14-1211, either D614 or D614G), containing the 2P and Furin cleavage site mutations^40^ with a mu-phosphatase signal peptide and a C-terminal Avi-8xHis-C-tag or C-terminal 8xHis-Avi-C-tag were expressed in Freestyle 293-F cells at 37°C and 8% CO_2_. Transfections were performed using 293fectin as a transfection reagent. Cell culture supernatant was collected after three days and purified over a 5 mL C-tag affinity matrix. Elution fractions were concentrated and injected on a Superose 6 Increase 10/300 GL column with 50 mM Tris-HCl pH 8.0 and 200 mM NaCl as running buffer.

SARS-CoV-2 HexaPro Spike protein for cryoEM analysis was produced in HEK293F cells grown in suspension using FreeStyle 293 expression medium (Life Technologies) at 37°C in a humidified 8% CO2 incubator rotating at 130 r.p.m. The cultures were transfected using PEI (9 μg/mL) with cells grown to a density of 2.5 million cells per mL and cultivated for 3 days. The supernatants were harvested and cells resuspended for another 3 days, yielding two harvests. Spike proteins were purified from clarified supernatants using a 5mL Cobalt affinity column (Cytiva, HiTrap TALON crude), concentrated and flash frozen in a buffer containing 20 mM Tris pH 8.0 and 150 mM NaCl prior to analysis.

Recombinant hACE2 for SPR (residues 19-615 from Uniprot Q9BYF1 with a C-terminal AviTag-10xHis-GGG-tag, and N-terminal signal peptide) was expressed in HEK293.sus using standard methods (ATUM Bio). Protein was purified via Ni Sepharose resin followed by isolation of the monomeric hACE2 by size exclusion chromatography using a Superdex 200 Increase 10/300 GL column pre-equilibrated with PBS.

### SPR binding assays

SPR binding measurements were performed using a Biacore T200 instrument with CM5 sensor chip covalently immobilized with StrepTactin XT to capture recombinant RBD proteins (data in Fig. 1a and Extended Data Fig. 5f). Running buffer was Cytiva HBS-EP+ (pH 7.4). All measurements were performed at 25°C. Fab (or hACE2) analyte concentrations were 11, 33, 100, and 300 nM, run as single-cycle kinetics. Double reference-subtracted data were fit to a 1:1 binding model using Biacore Evaluation or Biacore Insight software. K_D_ above 1 μM were determined from fits where Rmax was set as a constant based on results for higher affinity analytes binding to the same RBD at the same surface density. Data are representative of duplicate measurements.

To corroborate the SARS-CoV-2 RBD binding measurements, experiments were also performed in two additional formats, both with monovalent analytes (data in Extended Data Table 1): (1) Fab binding to SARS-CoV-2 spike ectodomain was measured using CM5 sensor chips immobilized with anti-AviTag pAb for capturing S, other experiment parameters same as above, and (2) RBD binding to IgG was measured using CM5 sensor chips immobilized with anti-human Fc pAb for capturing IgG, with RBD analyte concentrations of 3.1, 12.5, and 50 nM, other experiment parameters same as above. Fit results yield an “apparent K_D_” for the spike-binding experiments because the kinetics also reflect spike conformational dynamics. Spike ectodomain was D614G with C-terminal 8xHis-Avi-C-tag for all measurements except S2X58 binding was performed with D614 spike with C-terminal Avi-8xHis-C-tag.

### Deep mutational scanning mutant escape profiling

We used a previously described deep mutational scanning approach^3^ to comprehensively identify RBD mutations that escape binding of each antibody. This approach leverages duplicate RBD mutant libraries^28^, which contain virtually all of the 3,819 possible amino acid mutations in the background of the Wuhan-Hu-1 RBD sequence. Library variants were previously linked to short identifier barcode sequences and sorted to purge the library of variants that strongly decrease ACE2 binding affinity or expression of folded RBD^3^.

We first used an isogenic yeast strain expressing the unmutated SARS-CoV-2 RBD and flow cytometry to identify the EC90 of each antibody’s binding to yeast-displayed SARS-CoV-2 RBD. We then performed library selections as previously described^3,23^, labeling libraries with the EC90 concentration of antibody to standardize escape mutation sensitivity across selections. Briefly, libraries of yeast were induced for surface expression, washed, and labeled with the primary antibody for one hour at room temperature. Cells were washed, and secondarily labeled with 1:200 PE-conjugated goat anti-human-IgG antibody (Jackson ImmunoResearch 109-115-098) to label for bound antibody, and 1:100 FITC-conjugated chicken anti-Myc-tag (Immunology Consultants Lab, CYMC-45F) to label for RBD surface expression. We prepared controls for setting FACS selection gates by labeling yeast expressing the unmutated SARS-CoV-2 RBD with the same antibody concentration as library selections (1x), 100x reduced antibody concentration to illustrate the effect of mutations with 100x-reduced affinity, and 0 ng/mL antibody to illustrate complete loss of antibody binding. Representative selection gates are shown in Extended Data Fig. 2b. We sorted approximately 7.5e6 RBD+ cells per library on a BD FACSAria II, collecting yeast cells from the antibody-escape sort bin (fractions of library falling into antibody escape bin given in Extended Data Fig. 2c). Sorted cells were recovered overnight, plasmids were extracted from the pre-sort and antibody-escape populations, and variant-identifier barcode sequences were PCR amplified and sequenced on an Illumina HiSeq 2500^3,28^.

As previously described^3,23^, sequencing counts pre- and post-selection were used to estimate the “escape fraction” for each library variant, which reflects the fraction of yeast expressing a variant that fall into the antibody-escape FACS bin. Briefly, we used the dms_variants package (https://jbloomlab.github.io/dms_variants/, version 0.8.2) to process Illumina sequences into variant counts pre- and post-selection using the barcode/RBD variant lookup table from Starr et al.^28^. We then computed per-variant escape fractions as:

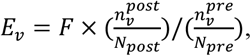

where *F* is the total fraction of the library that escapes antibody binding (Extended Data Fig. 2c), 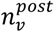 and 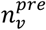 are the sequencing counts of variant *v* in the RBD library after and before FACS selection (with a pseudocount of 0.5 added to all counts), and *N*_*post*_ and *N*_*pre*_ are the total counts of all variants after and before FACS selection. We then applied computational filters to remove variants with low pre-sort sequencing counts or highly deleterious mutations that might cause artefactual antibody escape due to global unfolding or loss of expression of RBD on the cell surface. Specifically, we filtered out variants whose pre-selection sequencing counts were lower than the 99^th^ percentile counts of variants containing premature stop codons, which were largely purged by the prior sorts for RBD expressing and ACE2-binding RBD variants. We also removed variants with ACE2 binding scores < −2.35 or RBD expression scores < −1, and variants containing individual mutations with effects below these thresholds, using the variant- and mutation-level deep mutational scanning measurements from Starr et al.^28^. We also filtered out rare mutations with low coverage in the libraries, retaining mutations that were sampled on at least one single-mutant barcoded variant or at least two multiply-mutated variants in each replicate. Last, to decompose single-mutation escape fractions for each antibody, we implemented global epistasis models^41^ using the dms_variants package to estimate the effect of each individual amino acid mutation, exactly as described in ref. ^23^.

Antibody escape selections were conducted in full duplicate using independently generated and assayed SARS-CoV-2 mutant libraries (see correlations in Extended Data Fig. 2d,e). The reported escape fractions throughout the paper are the average across the two replicates, and these final per-mutation escape fractions are provided on GitHub: https://github.com/jbloomlab/SARS-CoV-2-RBD_MAP_Vir_mAbs/blob/main/results/supp_data/vir_antibodies_raw_data.csv. The interactive visualizations of antibody escape maps (https://jbloomlab.github.io/SARS-CoV-2-RBD_MAP_Vir_mAbs) were created using dms-view^42^.

### Sarbecovirus library binding assays

A curated set of all unique sarbecovirus RBD amino acid sequences was gathered, including the sarbecovirus RBD sequence set reported by Letko et al.^7^, along with additional unique RBD sequences among SARS-CoV-1 epidemic strains reported by Song et al.^43^, BtKY72^44^ and new sarbecovirus sequences RmYN02^45^, GD-Pangolin-CoV (consensus RBD reported in Fig. 3a of Lam et al.^46^), and GX-Pangolin-CoV^46^ (P2V, ambiguous nucleotide within codon 515 (SARS-CoV-2 spike numbering) resolved to retain F515, which is conserved in all other sarbecoviruses). A list of all RBDs and sequence accession numbers is available on GitHub: https://github.com/jbloomlab/SARSr-CoV_RBD_MAP/blob/main/data/RBD_accessions.csv

To define clades of sarbecovirus RBDs, an alignment of amino acid RBD sequences was generated using mafft^47^ with gap opening penalty 4.5 (alignment available on GitHub: https://github.com/jbloomlab/SARSr-CoV_RBD_MAP/blob/main/data/RBD_aa_aligned.fasta). The corresponding nucleotide sequence alignment was generated from the amino acid alignment using PAL2NAL^48^. The gene sequence phylogeny was inferred using RAxML version 8.2.12^49^, with the GTRGAMMA substitution model and a partition model with separate parameters for first, second, and third codon positions. The Hibecovirus RBD sequence Hp-Zhejiang2013 (Genbank: KF636752) was used as an outgroup for rooting of the sarbecovirus phylogeny.

All unique sarbecovirus RBD protein-coding sequences were ordered from IDT, Twist, and Genscript, and cloned into our yeast display vector^28^. Sequences were pooled and appended with downstream 16-nt barcode sequences according to the protocol described in Starr et al.^28^. Long read circular consensus sequences spanning the 16-nt barcode and RBD genotype were gathered on a PacBio Sequel v2.0 and processed exactly as described in Starr et al.^28^. This yielded a barcode:variant lookup table for the sarbecovirus RBD library analogous to that used for SARS-CoV-2 mutant libraries. This table is available on GitHub: https://github.com/jbloomlab/SARSr-CoV_RBD_MAP/blob/main/data/barcode_variant_table.csv.

The pooled sarbecovirus RBD library was labeled, sorted, and quantified as described for the SARS-CoV-2 mutant libraries above, except we only sorted ~1 million RBD+ cells due to the reduced library size. Sequencing and quantification of per-variant antibody escape was conducted as described above. Data for the HKU3-8 RBD is not shown, as this RBD did not express in our yeast-display platform. For several antibodies, we performed a secondary experiment, selecting the sarbecovirus RBD library with a more stringent “full escape” gate to select out only variants exhibiting complete loss of binding (Extended Data Fig. 2b,c).

For follow-up quantitative binding assays, select sarbecovirus RBDs were cloned into the yeast-display platform as isogenic stocks. Binding assays were conducted across a titration series of antibody in 96-well plates, and binding at each antibody concentration (geometric mean fluorescence intensity in the PE channel among RBD+ (FITC+) cells) was determined via flow cytometry, and fit to a four-parameter Hill curve to identify the EC50 (midpoint).

### Analysis of mutations in natural SARS-CoV-2 sequences

All spike sequences on GISAID^50^ as of March 4, 2021, were downloaded and aligned via mafft^47^. Sequences from non-human origins, sequences with gaps or ambiguous characters in the RBD, and sequences with more than 8 amino acid differences from the Wuhan-Hu-1 reference sequence (Genbank MN908947, residues N331-T531) were removed. We determined mutation frequencies compared to Wuhan-Hu-1 reference from this final alignment of 582,276 sequences. We acknowledge all contributors to the GISAID EpiCoV database for their sharing of sequence data. All contributors to GISAID EpiCoV listed at: https://github.com/jbloomlab/SARS-CoV-2-RBD_MAP_Vir_mAbs/blob/main/data/gisaid_hcov-19_acknowledgement_table_2021_03_04.pdf.

### Quantitative summary metrics of antibody properties

The relative epitope size of an antibody was calculated as the sum of per-mutant escape fractions that are at least five times the global median escape fraction (to minimize the impact of variation in background noise on the summation). For this summation, escape fractions were normalized to the maximum per-mutation escape fraction, to account for slight variation in the largest per-mutation escape fraction measured between selections.

The relative escapability of an antibody was calculated the same as relative epitope size, but each mutation was multiplied by two weighting factors scaled from 0 to 1 that reflect the impact of that mutation on ACE2-binding affinity and RBD expression as measured in our prior deep mutational scan^28^. The relationship between weighting factors and mutation effect on each property is shown in Extended Data Fig. 4b. Mutations with < –1 effect on either property are effectively zeroed out in the escapability summation. Mutations with effects between –1 and 0 have intermediate weights, and mutations with 0 or positive effects are assigned weight factors of 1.

Antibody susceptibility to escape by natural SARS-CoV-2 mutations was calculated as the summed GISAID frequencies of all escape mutations, where escape mutations (all labels in Extended Data Fig. 4) are defined as those with escape fraction greater than five times the median escape fraction as above. These summed natural escape frequences are tabulated in the plot headers in Extended Data Fig. 4a.

The summary breadth of an antibody was calculated from the sarbecovirus RBD library escape selection using the standard gating (Extended Data Fig. 5b), only. Although we have various follow-up binding data illustrating reduced affinity binding for some “escaped” sarbecovirus RBDs, these follow-up experiments were not conducted systematically for all antibody/RBD combinations, and therefore would bias breadth estimates. Breadth of binding was calculated as the frequency of all sarbecovirus RBDs that are bound with affinity within the FACS selection gating threshold, weighted by clade representation. Specifically, breadth was normalized to give equal representation to each of the four sarbecovirus clades to account for different depth of sampling. Within the SARS-CoV-1 clade, all human 02/03 strains and civet + human 03/04 strains were similarly downweighted to each represent 1/8 of the possible breadth within the SARS-CoV-1 clade (together with the six bat sarbecoviruses in this clade). As an example, breadth for S304 is calculated as [4/4 +([6/6]+[6/6]+5)/8 + 2/2 + 0/21]/4 = 0.72, based on the data shown in Extended Data Fig. 5b.

### Multidimensional scaling projection of antibody epitopes

Multidimensional scaling projection in Fig. 2 was performed using the Python scikit-learn package. We first computed the similarity and dissimilarity in the sites of escape between each pair of antibodies, exactly as described in Greaney et al.^3^, and performed metric multidimensional scaling with two components on the matrix of dissimilarities between all antibody pairs. Antibodies in this layout were colored with pie charts proportional to the total squared site-wise escape that falls into the labeled structural regions (RBM = residues 437 to 508, ACE2 contact defined as 4Å cutoff based on 6M0J crystal structure^51^, and core RBD otherwise). In this layout, we included all of our previously published antibodies for which we have performed escape mapping via this same approach. These antibodies and their citations include: S2X259, accompanying paper Tortorici et al. 2021; LY-CoV555^21^; COV2-2196 and COV2-2130^33^; REGN10933, REGN10987, and LY-CoV016^23^; and all other COV2 antibodies and CR3022^3^.

For Figures 2c-g, we colored the antibodies within this layout according to various antibody properties. When appropriate, we also colored these previously assayed antibodies, as described below. Extended Data Fig. 6 and the scatterplots in Fig. 2h-j show the relationships between properties for antibodies specifically in this study (and S2X259) for the most direct comparability.

Antibody neutralization potencies illustrated in Fig. 2c incorporate the authentic SARS-CoV-2 neutralization IC50s as reported in this study (Fig. 1a), together with the live SARS-CoV-2 neuralization IC50s for the COV2 antibodies reported by Zost et al.^10^. We acknowledge that it is imperfect to compare neutralization potencies reported from different labs on different antibody batches, though in this case, both sets are indeed neutralization potencies with authentic virus. We therefore do not directly compare these two sets of measurements in a quantitative manner, but we do note that their joint inclusion in Figure 2C supports the dichotomy between neutralization potency of core RBD versus RBM antibodies which is supported by either neutralization panel alone.

Sarbecovirus breadth illustrated in Fig. 2d incorporates the pan-sarbecovirus breadth measurements reported in the current study together with more limited breadth measurements for antibodies reported in our prior publications. These previously published experiments determined binding within a more restricted sarbecovirus RBD set present in our libraries (SARS-CoV-2, RaTG13, GD-Pangolin, SARS-CoV-1 [Urbani], LYRa11, and WIV1). We calculated breadth from this incomplete sarbecovirus sequence set for comparison, but note that these antibodies are limited to a relative breadth of 0.5 because no RBDs from the Africa/Europe or non-ACE2-utilizing Asia clades were included. However, as with neutralization, inclusion of these antibodies nonetheless emphasizes the core RBD/RBM dichotomy in sarbecovirus breadth established by our primary panel.

For illustrations of epitope size and escapability in Figs. 2e-g, we calculated these quantities for our previously profiled antibodies as described above. We excluded the antibodies profiled in Greaney et al.^3^, as these assays were performed on a prior version of our SARS-CoV-2 mutant library that exhibited different quantitative features of absolute escape, complicating its quantitative comparison to extent of escape for antibodies profiled in this and our other studies, which all use the same library.

Structural mappings around the perimeter of Fig. 2a were created by mapping total site-wise escape to the b-factor column of PDB structures. Footprints were defined as residues within a 5Å cutoff of antibody heavy atoms. Structures used were those described in this paper, or previously published structures: ACE2-bound RBD (6M0J)^51^, CR3022-bound RBD (6W41)^52^, REGN10987- and REGN10933-bound RBD (6XDG)^14^, CB6-(LY-CoV016) bound RBD (7C01)^16^, and S304, S309, and S2H14-bound RBD (7JX3)^17^.

### RBD ELISA

96 half area well-plates (Corning 3690) were coated over-night at 4°C with 25 μL of sarbecoviruses RBD proteins at 5 μg/mL in PBS pH 7.2. Plates were blocked with PBS 1% BSA (Sigma-Aldrich, A3059) and subsequently incubated with mAb serial dilutions for 1 h at room temperature. After 4 washing steps with PBS 0.05% Tween 20 (PBS-T) (Sigma-Aldrich, 93773), goat anti-human IgG secondary antibody (Southern Biotech, 2040-04) was added and incubated for 1 h at room temperature. Plates were then washed 4 times with PBS-T and 4-NitroPhenyl phosphate (pNPP, Sigma-Aldrich, 71768) substrate added. After 30 min incubation, absorbance at 405 nm was measured by a plate reader (Biotek) and data plotted using Prism GraphPad.

### Binding to cell surface expressed sarbecovirus S proteins by flow cytometry

ExpiCHO-S cells were seeded at 6 × 10^6^ cells cells/mL in a volume of 5 mL in a 50 mL bioreactor. Spike coding plasmids were diluted in cold OptiPRO SFM, mixed with ExpiFectamine CHO Reagent (Life Technologies) and added to the cells. Transfected cells were then incubated at 37°C with 8% CO_2_ with an orbital shaking speed of 120 RPM (orbital diameter of 25 mm) for 42 hours. Transiently transfected ExpiCHO cells were harvested and washed two times in wash buffer (PBS 1% BSA, 2 mM EDTA). Cells were counted and distributed into round bottom 96-well plates (Corning) and incubated with 10 μg/mL S2H97, S2X35 or S309 mAb. Alexa Fluor647-labelled Goat Anti-Human IgG secondary Ab (Jackson ImmunoResearch 109-607-003) was prepared at 1.5 mg/mL added onto cells after two washing steps. Cells were then washed twice and resuspended in wash buffer for data acquisition at ZE5 cytometer (Biorad).

### Selection of VSV-SARS-CoV-2 monoclonal antibody escape mutants (MARMS)

VSV-SARS-CoV-2 chimera was used to select for SARS-CoV-2 S monoclonal antibody resistant mutants (MARMS) as previously described^1,53^. Briefly, MARMS were recovered by plaque isolation on Vero E6 cells with the indicated mAb in the overlay. The concentration of mAb in the overlay was determined by neutralization assays at a multiplicity of infection (MOI) of 100. Escape clones were plaque-purified on Vero cells in the presence of mAb, and plaques in agarose plugs were amplified on MA104 cells with the mAb present in the medium. Viral stocks were amplified on MA104 cells at an MOI of 0.01 in Medium 199 containing 2% FBS and 20 mM HEPES pH 7.7 (Millipore Sigma) at 34°C. Viral supernatants were harvested upon extensive cytopathic effect and clarified of cell debris by centrifugation at 1,000 × g for 5 min. Aliquots were maintained at −80°C. Viral RNA was extracted from VSV-SARS-CoV-2 mutant viruses using RNeasy Mini kit (Qiagen), and S was amplified using OneStep RT-PCR Kit (Qiagen). The mutations were identified by Sanger sequencing (GENEWIZ). Their resistance was verified by subsequent virus infection in the presence or absence of antibody. Briefly, Vero cells were seeded into 12 well plates for overnight. The virus was serially diluted using DMEM and cells were infected at 37°C for 1 h. Cells were cultured with an agarose overlay in the presence or absence of mAb at 34°C for 2 days. Plates were scanned on a biomolecular imager and expression of eGFP is shown at 48 hours post-infection.

### Crystallization, data collection, structure determination, and analysis

To form RBD-Fab complexes for crystallization, SARS-CoV-2 RBD was mixed with a 1.3-fold molar excess of each Fab and incubated on ice for 20-60 min. Complexes were purified on a Superdex 200 Increase 10/300 GL column preequilibrated with 20 mM Tris-HCl pH 7.5 and 150 mM NaCl. Crystals of the RBD-Fab complexes were obtained at 20°C by sitting drop vapor diffusion.

For the SARS-CoV-2 RBD-S2X35-S309 complex, a total of 200 nL complex at 5.4 mg/mL was mixed with 100 nL mother liquor solution containing 1.85 M Ammonium Sulfate, 0.1 M Tris pH 8.17, 0.8% (w/v) polyvinyl alcohol, 1% (v/v) 1-propanol, and 0.01 M HEPES pH 7. Crystals were flash frozen in liquid nitrogen using the mother liquor solution supplemented with 20% glycerol for cryoprotection. Data were collected at Beamline 9-2 of the Stanford Synchrotron Radiation Lightsource and processed with the XDS software package (Kabsch, 2010) to 1.83 Å in space group C 2 2 2. The RBD-S2X35-S309 Fab complex structure was solved by molecular replacement using phaser^54^ from a starting model consisting of RBD-S309 Fab (PDB ID: 7JX3) and a homology model for the S2X35 Fab built using the Molecular Operating Environment (MOE) software package from the Chemical Computing Group (https://www.chemcomp.com).

For the SARS-CoV-2-RBD-S2H97-S2X259 Fab complex, 200 nL complex at 5.7 mg/mL were mixed with 200 nL mother liquor solution containing 0.12 M Monosaccharides, 20% (v/v) Ethylene glycol, 10% (w/v) PEG 8000, 0.1 M Tris (base)/bicine pH 8.5, 0.02 M Sodium chloride, 0.01 M MES pH 6 and 3% (v/v) Jeffamine ED-2003. Crystals were flash frozen in liquid nitrogen. Data were collected at Beamline 9-2 of the Stanford Synchrotron Radiation Lightsource facility in Stanford, CA. Data were processed with the XDS software package (Kabsch, 2010) for a final dataset of 2.65 Å in space group P 21. The RBD-S2H97-S2X259 Fab complex structure was solved my molecular replacement using phaser from a starting model consisting of SARS-CoV-2 RBD (PDB ID: 7JX3) and homology models for the S2H97 and S2X259 Fabs built using the MOE software package.

For the SARS-CoV-2-RBD-S2E12-S304-S309 Fab complex, 200 nL complex at 4.5 mg/mL were mixed with 100 nL of 0.09 M Phosphate/Citrate pH 5.5, 27% (v/v) PEG Smear Low, 4% (v/v) Polypropylene glycol 400 and 100 nL of 0.02 M Imidazole pH 7 or 0.09 M Phosphate/Citrate pH 5.5, 27% (v/v) PEG Smear Low, 0.01 M Potassium/sodium phosphate pH 7, 1% (v/v) PPGBA 230 and 1.5% (v/v) PPGBA 400. Crystals were flash frozen in liquid nitrogen. Data were collected at the Molecular Biology Consortium beamline 4.2.2 at the Advanced Light Source synchrotron facility in Berkeley, CA. Datasets from two crystals from the two conditions were individually processed and then merged with the XDS software package^55^ for a final dataset of 2.93 Å in space group I 41 2 2. The RBD-S2E12-S304-S309 Fab complex structure was solved my molecular replacement using phaser from starting models consisting of RBD-S304-S309 Fab (PDB ID: 7JX3) and S2E12 (PDB ID: 7K3Q).

For all structures, several subsequent rounds of model building and refinement were performed using Coot^56^, ISOLDE^57^, Refmac5^58^, and MOE (https://www.chemcomp.com), to arrive at the final models. For all complexes, epitopes on the RBD protein were determined by identifying all RBD residues within a 5.0 Å distance from any Fab atoms. The analysis was performed using the MOE software package and the results were manually confirmed.

### Cryo-electron microscopy

SARS-CoV-2 HexaPro S^59^ at 1.2 mg/mL was incubated with 1.2 fold molar excess of recombinantly purified S2D106 or S2H97 at 4°C before application onto a freshly glow discharged 2.0/2.0 UltrAuFoil grid (200 mesh). Plunge freezing used a vitrobot MarkIV (Thermo Fisher Scientific) using a blot force of 0 and 6.5 second blot time at 100% humidity and 23°C.

For the S/S2D106 data set, Data were acquired using an FEI Titan Krios transmission electron microscope operated at 300 kV and equipped with a Gatan K2 Summit direct detector and Gatan Quantum GIF energy filter, operated in zero-loss mode with a slit width of 20 eV. Automated data collection was carried out using Leginon^60^at a nominal magnification of 130,000x with a pixel size of 0.525 Å. The dose rate was adjusted to 8 counts/pixel/s, and each movie was acquired in super-resolution mode fractionated in 50 frames of 200 ms. 2,166 micrographs were collected with a defocus range between −0.5 and −2.5 μm. Movie frame alignment, estimation of the microscope contrast-transfer function parameters, particle picking, and extraction were carried out using Warp^61^. Particle images were extracted with a box size of 800 binned to 400 pixels^2 yielding a pixel size of 1.05 Å.

For the S/S2H97 data set, data were acquired on an FEI Glacios transmission electron microscope operated at 200 kV equipped with a Gatan K2 Summit direct detector. Automated data collection was carried out using Leginon^60^at a nominal magnification of 36,000x with a pixel size of 1.16 Å. The dose rate was adjusted to 8 counts/pixel/s, and each movie was acquired in counting mode fractionated in 50 frames of 200 ms. 3,138 micrographs were collected in a single session with a defocus range comprised between −0.5 and −3.0 μm. Preprocessing was performed using Warp^[Citation error]^ and particle images were extracted with a box size of 400 pixels^2.

For the S/S2D106 and S/S2H97 datasets, two rounds of reference-free 2D classification were performed using CryoSPARC to select well-defined particle images^62^. These selected particles were subjected to two rounds of 3D classification with 50 iterations each (angular sampling 7.5° for 25 iterations and 1.8° with local search for 25 iterations), using our previously reported closed SARS-CoV-2 S structure as initial model^40^ (PDB 6VXX) in Relion^63^. 3D refinements were carried out using non-uniform refinement^64^ along with per-particle defocus refinement in CryoSPARC. Selected particle images were subjected to the Bayesian polishing procedure^65^ implemented in Relion3.0 before performing another round of non-uniform refinement in CryoSPARC followed by per-particle defocus refinement and again non-uniform refinement.

To further improve the density of the S2D106 Fab, the particles were then subjected to focus 3D classification without refining angles and shifts using a soft mask on RBD and Fab region with a tau value of 60 in Relion. Particles belonging to classes with the best resolved local density were selected and subject to local refinement using CryoSPARC. Local resolution estimation, filtering, and sharpening were carried out using CryoSPARC. Reported resolutions are based on the gold-standard Fourier shell correlation (FSC) of 0.143 criterion and Fourier shell correlation curves were corrected for the effects of soft masking by high-resolution noise substitution^66^. UCSF Chimera^67^ and Coot^68^ were used to fit atomic models into the cryoEM maps. Spike-RBD/S2D106 Fab model was refined and relaxed using Rosetta using sharpened and unsharpened maps^69^.

### Molecular dynamics simulations

Full details of molecular dynamics workflow and analysis are available on GitHub: https://github.com/choderalab/rbd-ab-contact-analysis. The WT RBD:S309 complex was constructed from PDB ID 7JX3 (Chains A, B, and R). 7JX3 was first refined using ISOLDE^57^ to better fit the experimental electron density using detailed manual inspection. Refinement included adjusting several rotamers, flipping several peptide bonds, fixing several weakly resolved waters, and building in a missing four-residue-long loop. Though the N343 glycan N-Acetylglucosamine (NAG) was present in 7JX3, ISOLDE was used to construct a complex glycan at N343. The full glycosylation pattern was determined from Shajahan et al.^70^ and Watanabe et al.^71^ The glycan structure used for N343 (FA2G2) corresponds to the most stable conformer obtained from multi microsecond molecular dynamics (MD) simulations of cumulative sampling.^72^ The base NAG residue in FA2G2 was aligned to the corresponding NAG stub in the RBD:S309 model and any resulting clashes were refined in ISOLDE. The fucose included in the N343 glycan was modeled as the D-enantiomer. Mutant RBD:S309 complexes were constructed from the fully glycosylated WT RBD:S309 complex using PyMOL 2.3.5 (Schrödinger, LLC).

The refined glycosylated RBD:S309 complexes were prepared for simulation using tleap from AmberTools20^73^. All relevant disulfide bridges and covalent connections in glycan structures were specified. The glycosylated proteins were parameterized with the Amber ff14SB^74^ and GLYCAM_06j-1^75^ force fields. The systems were solvated using the TIP3P rigid water model^76^ in a truncated octahedral box with 2.2 nm solvent padding on all sides. The solvent box’s shape and size were chosen to prevent the protein complex from interacting with its periodic image. The solvated systems were then neutralized with 0.15 M NaCl using the Li/Merz ion parameters of monovalent ions for the TIP3P water model (12-6 normal usage set)^77^. Virtual bonds were added across chains that should be imaged together to aid the post-processing of trajectories.

The systems were energy-minimized with an energy tolerance of 10 kJ mol^−1^ and equilibrated using the OpenMMTools 0.20.0 (https://github.com/choderalab/openmmtools) BAOAB Langevin integrator^78^ for 20 ns in the NPT (p=1 atm, T = 310 K) ensemble with a timestep of 4.0 femtoseconds, a collision rate of 1.0 picoseconds^-1^, and a relative constraint tolerance of 1 ✕ 10^−5^. Hydrogen atom masses were set to 4.0 amu by transferring mass from connected heavy atoms, bonds to hydrogen were constrained, and center of mass motion was not removed. Pressure was controlled by a molecular-scaling Monte Carlo barostat with an update interval of 25 steps. Non-bonded interactions were treated with the Particle Mesh Ewald method^79^ using a real-space cutoff of 1.0 nm and the OpenMM default relative error tolerance of 0.0005, with grid spacing selected automatically. The simulations were subsequently packaged to seed for production simulation on Folding@home^80,81^. Default parameters were used unless noted otherwise.

The equilibrated structures were used to initiate parallel distributed MD simulations on Folding@home^80,81^. Simulations were run with OpenMM 7.4.2 (compiled into Folding@home core22 0.0.13). Production simulations used the same Langevin integrator as the NPT equilibration described above. In total, 3000 independent MD simulations were generated on Folding@home. Conformational snapshots (frames) were stored at an interval of 1 ns/frame for subsequent analysis. The resulting final dataset contained 3000 trajectories with 189 μs of aggregate simulation time. This amount of simulation time corresponds to ~20 GPU-years on an NVIDIA GeForce GTX 1080. This trajectory dataset (without solvent) will be available at the MolSSI COVID-19 Molecular Structure and Therapeutics Hub.

Total simulation times of ~42, 56, and 91 μs were used for analysis of WT, P337A, and P337L RBD:S309 systems, respectively. All trajectories had solute structures aligned to their first frame and centered using MDTraj^82^. RBD / S309 residues were considered to be at the RBD:S309 interface if they were within 10 Å of any S309 / RBD residue (with the exception of RBD N343 glycans, where all glycan residues were considered). The minimum distance of heavy atoms between every pair of interface residues was computed for every frame (1 ns) using MDAnalysis^83,84^. A close contact was counted if the minimum distance between a residue pair was below 3.5 Å. The contribution of each RBD residue to close contacts was calculated as a percentage by summation of the number of close contacts for a particular RBD residue and normalizing by the total number of close contact interactions over all frames of each simulation.

## Materials and Data Availability

- The SARS-CoV-2 RBD mutant libraries (#1000000172) and unmutated parental plasmid (#166782) are available on Addgene
- Other materials generated in this study will be made available on request and may require a material transfer agreement
- Raw Illumina sequencing data from deep mutational scanning experiments are available on NCBI SRA, BioSample SAMN18315604 (SARS-CoV-2 mutant selection data) and BioSample SAMN18316011 (sarbecovirus RBD selection data).
- PacBio sequencing data used to link N16 barcodes to sarbecovirus RBD variant are available on NCBI SRA, BioSample SAMN18316101.
- Complete table of deep mutational scanning antibody escape fractions is provided on GitHub: https://github.com/jbloomlab/SARS-CoV-2-RBD_MAP_Vir_mAbs/blob/main/results/supp_data/all_antibodies_raw_data.csv. This table includes both antibodies first described in this study (Fig. 1b,c), and all other antibody selections that were re-processed to generate Fig. 2a.
- The X-ray structure data and model has been deposited with accession code PDB XXX for RBD-S2X35-S309, PDB 7M7W for RBD-S2H97-S2X259 and PDB XXX for RBD-S2E12-S304-S309.
- The raw and processed molecular dynamics trajectory data will be available at the MolSSI COVID-19 Molecular Structure and Therapeutics Hub

## Code Availability

- Repository containing all code, analysis, and summary notebooks for the analysis of the SARS-CoV-2 deep mutational scanning escape selections available on GitHub: https://github.com/jbloomlab/SARS-CoV-2-RBD_MAP_Vir_mAbs
- Repository containing code and analysis of the sarbecovirus RBD library binding experiments available on GitHub: https://github.com/jbloomlab/SARSr-CoV_RBD_MAP
- Repository containing code and analysis of molecular dynamics simulations is available on GitHub: https://github.com/choderalab/rbd-ab-contact-analysis

---

**Extended Data Fig. 1.**
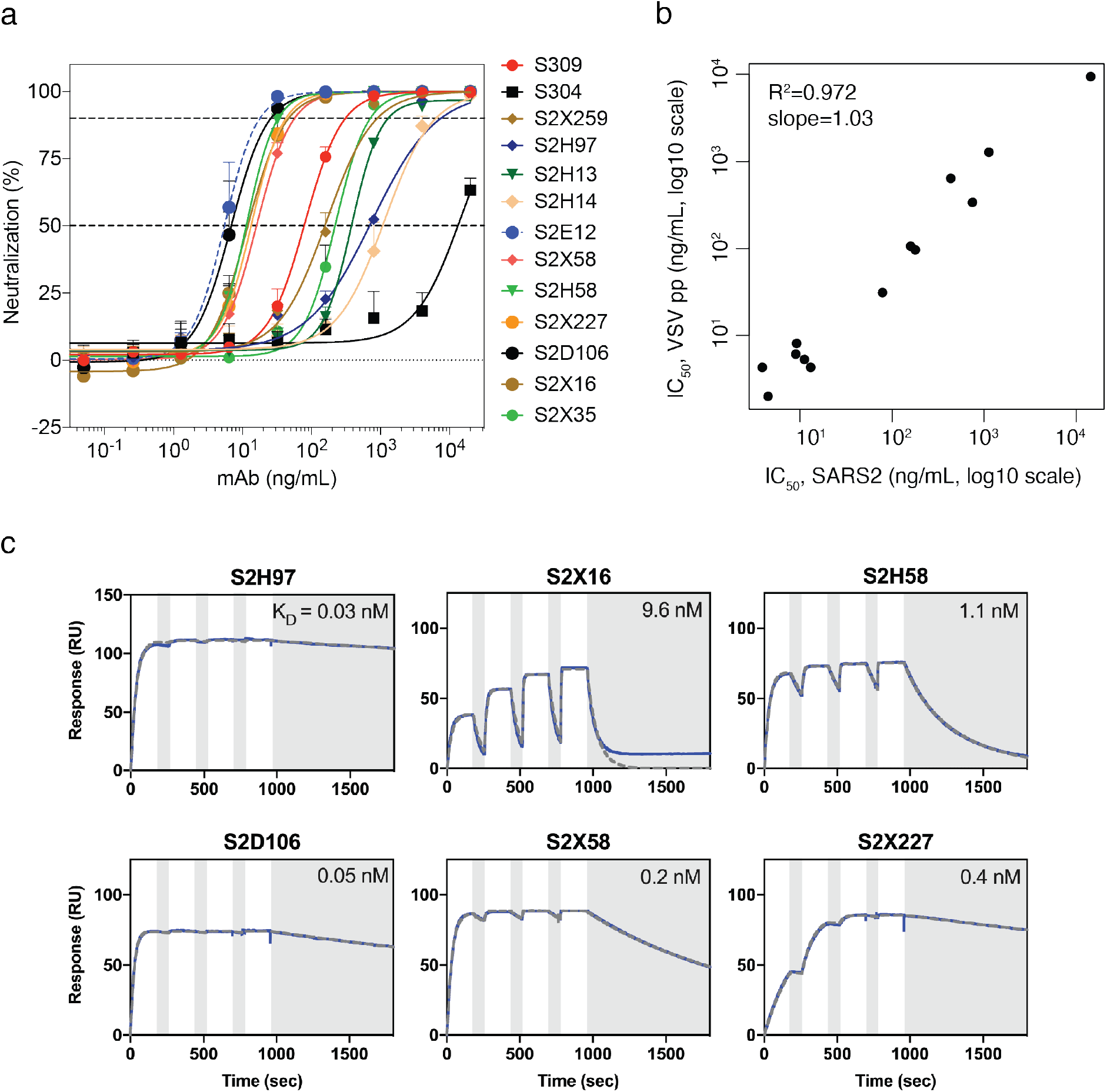
Antibody neutralization and binding data. **a**, Neutralization of authentic SARS-CoV-2 (SARS-CoV-2-Nluc) by 13 antibodies. Shown are representative live virus neutralization plots. Symbols are means ± SD of triplicates. Dashed lines indicate IC50 and IC90 values. **b**, Correlation in antibody neutralization IC_50_ as determined in spike-pseudotyped VSV particles (*n* = 3 to 8) versus authentic SARS-CoV-2 (*n* = 3). **c**, SPR analysis of Fab fragments of the six newly described antibodies binding to the SARS-CoV-2 RBD, summarized in Fig. 1a. White and gray stripes indicate association and dissociation phases, respectively.

**Extended Data Fig. 2.**
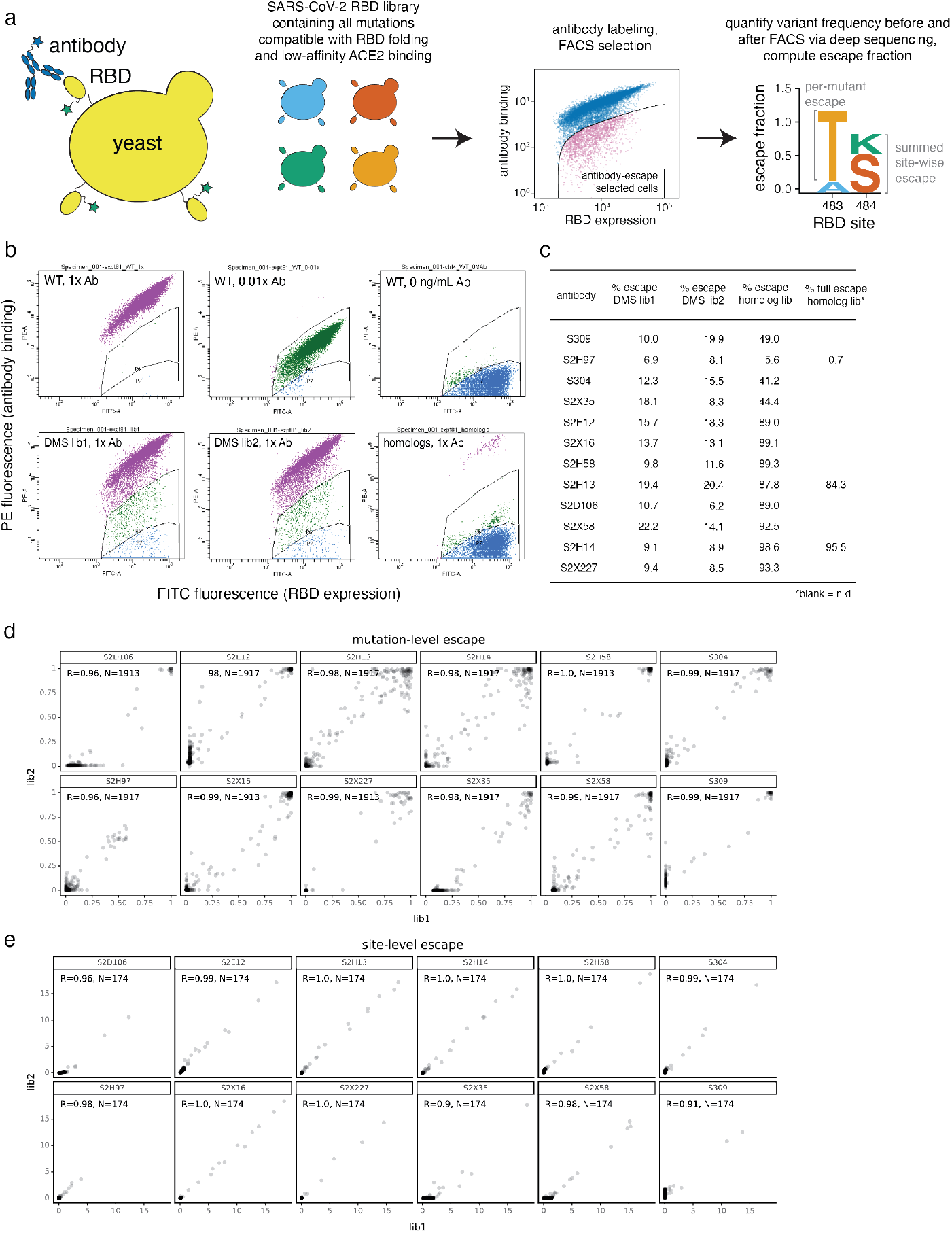
Yeast-display deep mutational scanning to comprehensively map mutations that escape antibody binding. **a**, Scheme of the deep mutational scanning assay. Conformationally intact RBD is expressed on the surface of yeast, where RBD expression and antibody binding is detectable via fluorescent labeling. We previously constructed mutant libraries containing virtually all of the 3,819 possible amino acid mutations in the SARS-CoV-2 RBD^28^ and sorted the library to eliminate mutations that destabilize the RBD or reduce ACE2-binding affinity by more than two orders of magnitude^3^. To identify mutations that escape antibody binding, we incubate the library with a sub-saturating antibody concentration (EC90 as determined by pilot yeast-displayed SARS-CoV-2 RBD binding assays) and use fluorescence-activated cell sorting (FACS) to isolate yeast cells expressing RBD mutants with reduced antibody binding. We use deep sequencing to quantify mutant frequencies before and after FACS selection, enabling calculation of the “escape fraction” of each amino acid mutation, which reflects the fraction of cells carrying that mutation that fall into the antibody-escape bin. Mutation escape fractions are represented in logoplots, where the height of a letter reflects the extent of escape from antibody binding. In this work, we created analogous libraries of yeast expressing all sarbecovirus RBDs, which we select via the same approach of FACS and deep sequencing to calculate the escape fraction of each sarbecovirus RBD. **b**, Representative selection gates, after gating for single cells expressing RBD as in Greaney et al.^3^. Yeast expressing the SARS-CoV-2 RBD (top panels) are labeled at 1x, 0.01x and no antibody to guide selection gates. Mutant RBDs that reduce binding (green, gate drawn to capture 0.01x WT control) are sorted and sequenced for calculation of mutant escape fractions. This same gate was used to quantify escape within libraries of yeast expressing all sarbecovirus RBD homologs. For several antibodies, we also selected the sarbecovirus RBD library with a more stringent “full escape” gate (blue, gate drawn to capture 0 ng/mL WT control), to enable differentiation between homologs that partially or completely escape antibody binding. **c**, Fraction of library cells falling into escape bins for each antibody selection. **d**, Correlation in per-mutation escape fractions for duplicate libraries that were independently generated and assayed. **e**, Correlation in per-site escape (sum of per-mutation escape fractions) for duplicate libraries.

**Extended Data Figure 3.**
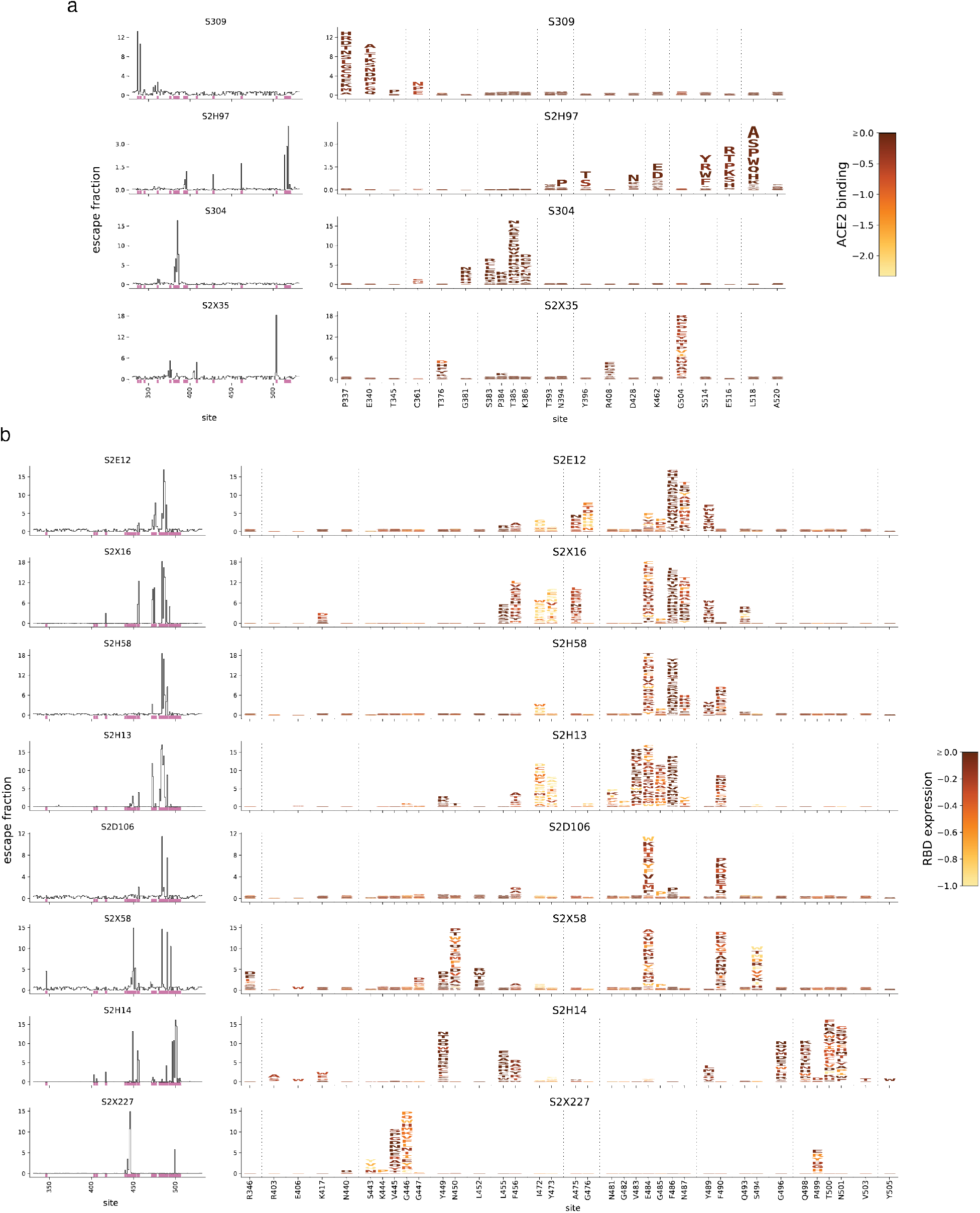
Expanded antibody escape profiles. **a**,**b**, Complete maps of escape, with mutations colored by their effects on the opposite RBD property illustrated in Fig. 1b,c. Line plots to the left of each logoplot illustrate the total escape at each RBD site. Pink bars illustrate sites shown in logoplots at right. Sites in logoplots are more expansive than Fig. 1b,c, as they reflect a more sensitive threshold of site-wise escape for logoplot visualization due to space constraints in Fig. 1b,c.

**Extended Data Fig. 4.**
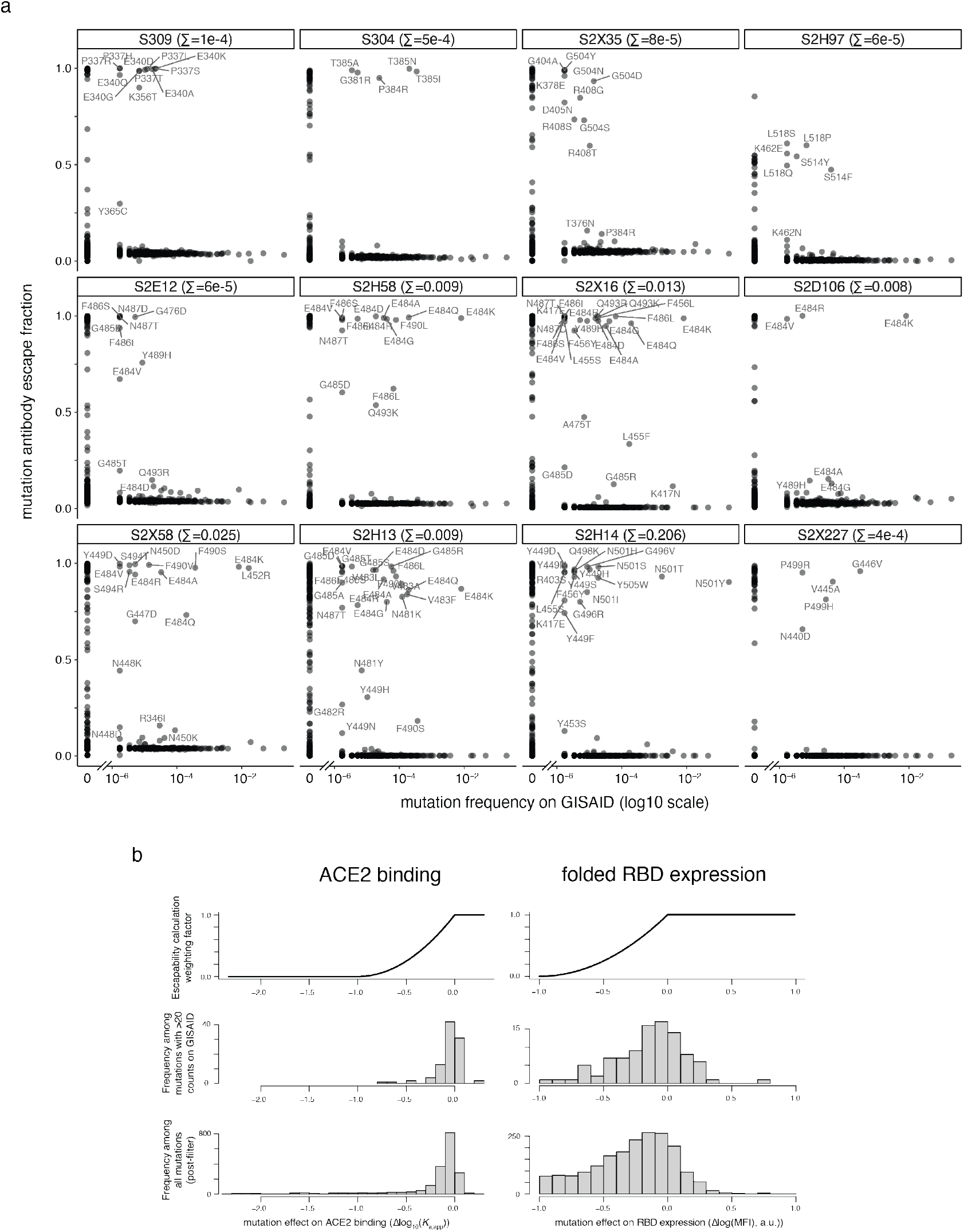
Antibody escapability in natural SARS-CoV-2 mutants and from deep mutational scanning measurements. **a**, For each antibody, scatterplots illustrate the degree to which a mutation escapes antibody binding (escape fraction, y-axis) versus its frequency among 582,276 high-quality human-derived SARS-CoV-2 sequences present on GISAID as of March 4, 2021. Large escape mutations (>5x global median escape fraction) for each antibody with non-zero mutant frequencies are labeled. The sum in each plot label gives the sum of mutant frequencies for all labeled mutations, corresponding to the natural SARS-CoV-2 mutant escape frequency for antibodies shown in Figs. 2g,j. **b**, To calculate antibody escapability (Fig. 1b,c), mutation escape fractions were weighted by their deleterious consequences for ACE2 binding or RBD expression. Top plots show the weighting factor (y-axis) for mutation effects on ACE2 binding (left) and RBD expression (right). This scaling weight factor was multiplied by the mutation escape fraction in the summation to calculate antibody escapability, as described in the Methods. For contextualizing this weighting penalization, histograms show the distribution of mutation effects on ACE2 binding (left) and RBD expression (right) for all mutations that pass our computational filtering steps (bottom), and mutations that are found with at least 20 sequence counts on GISAID (middle).

**Extended Data Fig. 5.**
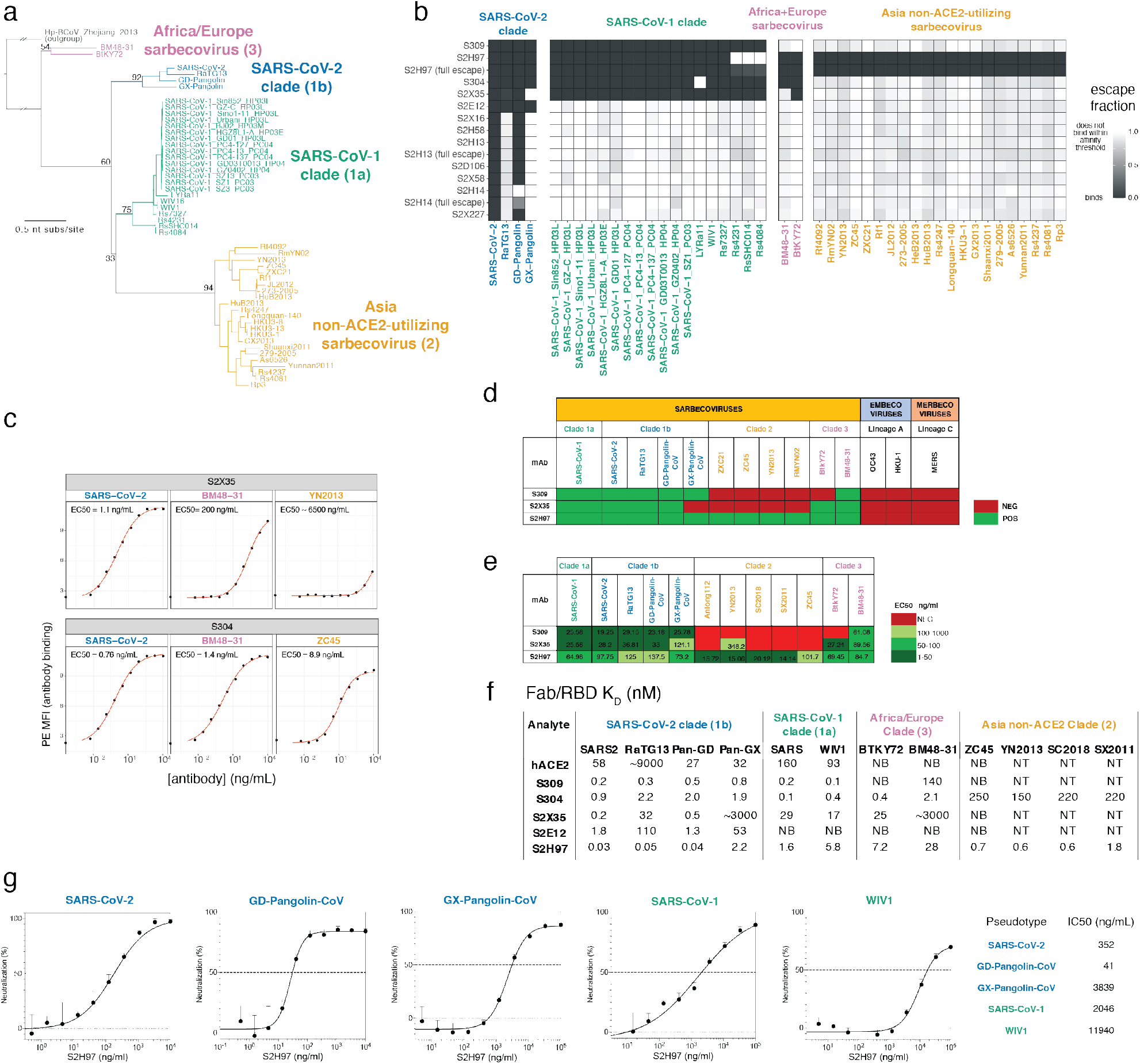
Breadth of antibody binding across sarbecoviruses. **a**, Phylogenetic relationship of sarbecovirus RBDs inferred from aligned RBD nucleotide sequences, with the four clades of sarbecovirus RBD labeled in separate colors used throughout text. Node support values are rapid bootstrap support values, illustrating substantial ambiguity in the exact relationship between the three clades of sarbecovirus in Asia. **b**, A yeast-display library containing all sarbecovirus RBDs was assayed for antibody escape analogous to SARS-CoV-2 mutant selections, as shown in Extended Data Fig. 2. Heatmaps show escape (white) versus binding (black) within the affinity threshold of the FACS escape bins (Extended Data Fig. 2b). For S2H97, S2H13, and S2H14, we repeated selections with a more stringent “full escape” bin (0 ng/mL WT control, Extended Data Fig. 2b,c), enabling differentiation of RBDs with intermediate binding (e.g., S2H97/RsSHC014) versus complete loss of binding. **c-e**, Because the high-throughput assay in (**b**) yields binary measures of binding versus escape at a set threshold determined by FACS gate selection, we performed follow-up quantitative binding assays for select antibody/RBD combinations including flow cytometry detection of antibody binding to isogenic yeast-displayed RBD variants (**c**), flow cytometry detection of antibody binding to mammalian-surface displayed spikes (**d**), and ELISA using purified RBD proteins (**e**). These experiments validate the affinity thresholds of “escape” illustrated in (**b**) while adding additional context to interactions that are still present but with reduced binding strength. **f**, Binding of cross-reactive antibodies (Fab) and human ACE2 to select sarbecovirus RBDs was determined via SPR. NB, no binding; NT, not tested. **g**, S2H97 neutralization of VSV pseudotyped with select sarbecovirus spikes, with entry measured in VeroE6 (SARS-CoV-2, GD-Pangolin-CoV, and SARS-CoV-1) or ACE2-transduced BHK-21 cells (GX-Pangolin-CoV and WIV1). Curves are representative of at least two independent experiments. Error bars represent standard deviation from three technical replicates from one representative experiment.

**Extended Data Fig. 6.**
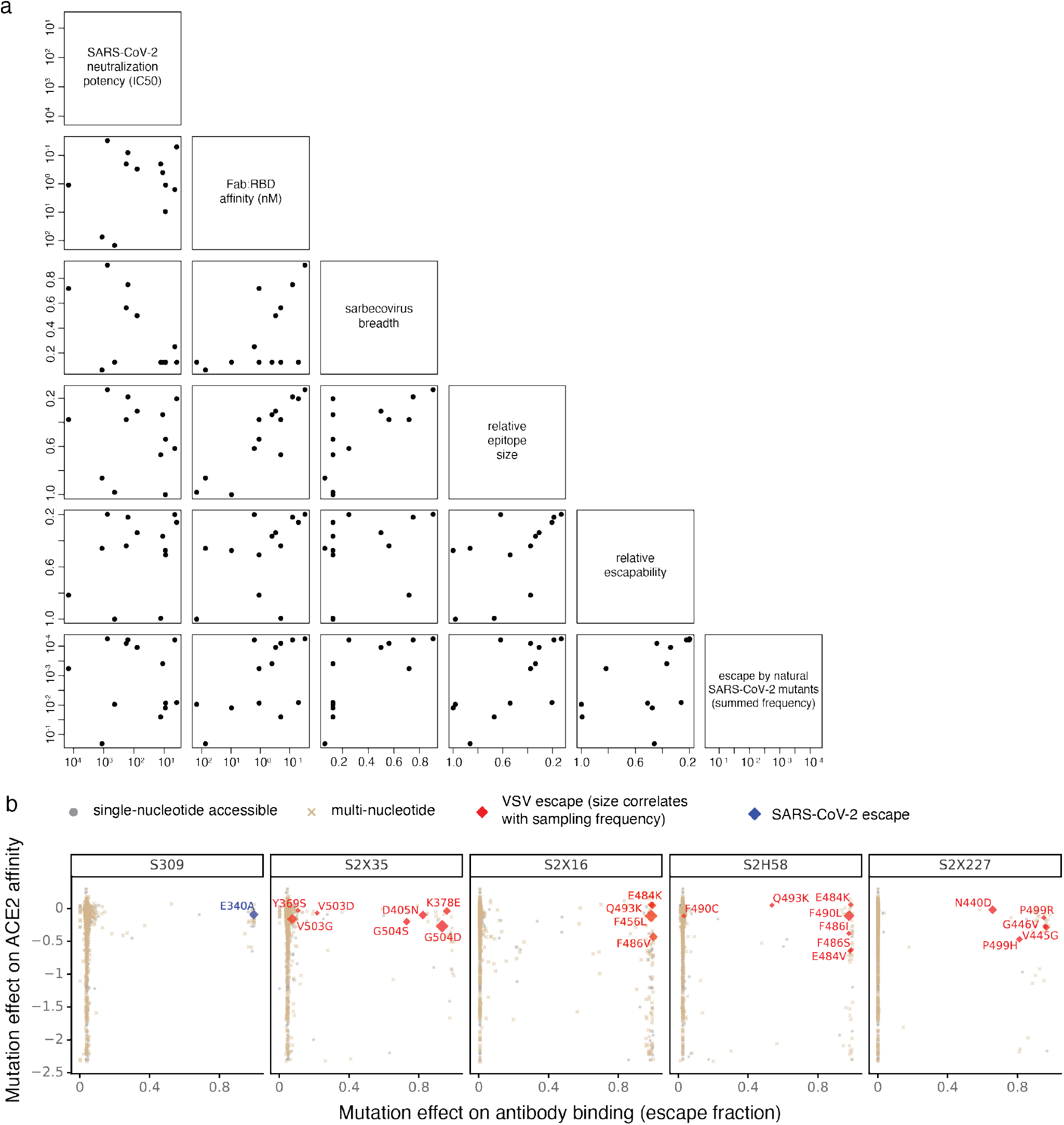
Antibody escapability and its relationship to other properties. **a**, Pairwise scatterplots between all antibody properties discussed in the main text. Select scatterplots from this panel are shown in Figs. 2h-j. Details of each property described in Methods. All axes are oriented such that moving to up on the y-axis and right on the x-axis corresponds to moving in the “preferred” direction for an antibody property (lower neutralization IC50, lower *K*_D_, higher breadth, narrower epitope size, lower escapability, lower total frequency of SARS-CoV-2 escape mutants among sequences on GISAID). **b**, Additional viral escape selections, as in Fig. 3a. For S309, we also include the E340A mutation identified in selections of authentic SARS-CoV-2 by Cathcart et al.^24^.

**Extended Data Fig. 7.**
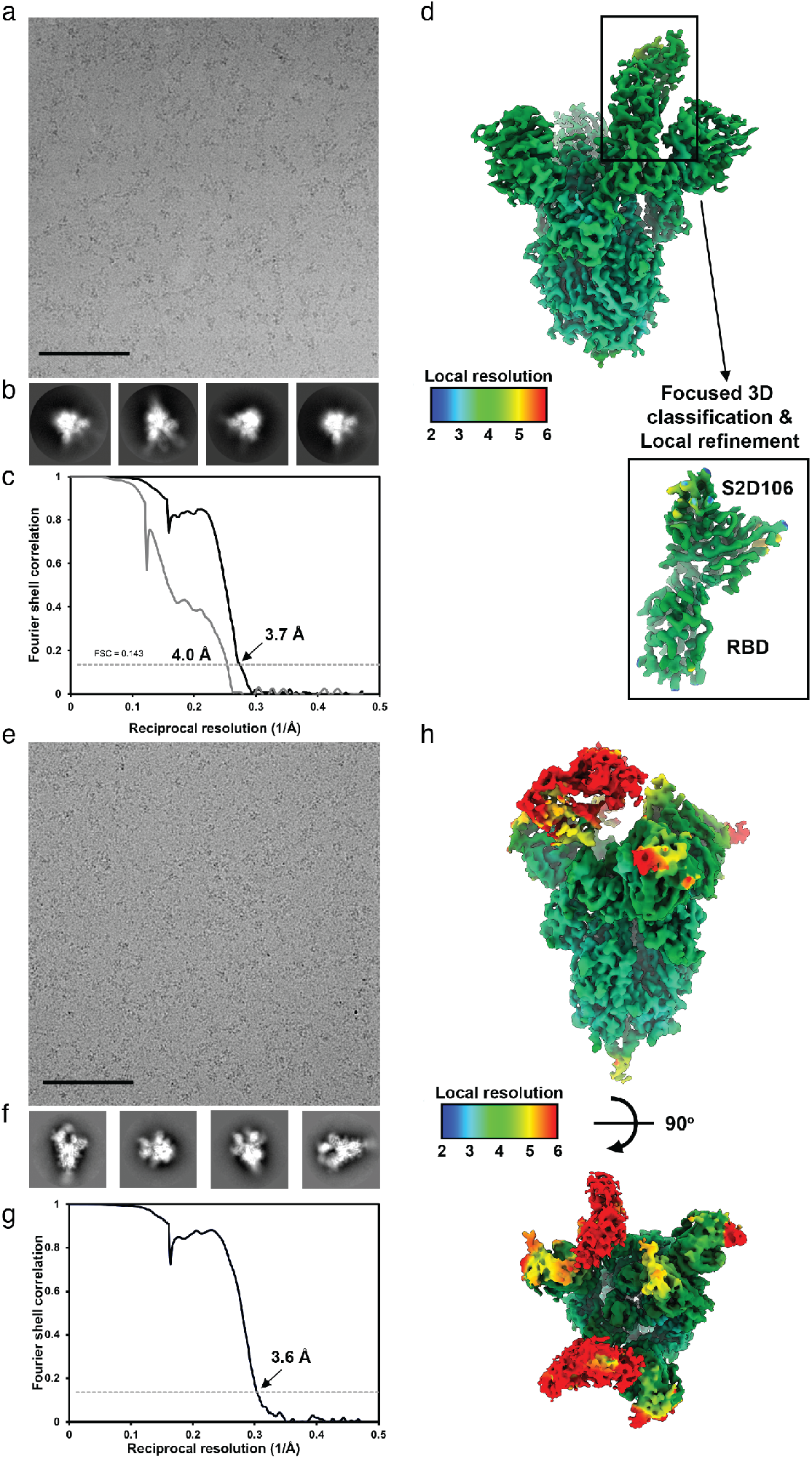
Data collection and processing of the S/S2D106 and S/S2H97 complex cryoEM datasets. **a,b**, Representative electron micrograph and 2D class averages of SARS-CoV-2 S in complex with the S2D106 Fab embedded in vitreous ice. Scale bar: 400 Å. **c**, Gold-standard Fourier shell correlation curves for the S2D106-bound SARS-CoV-2 S trimer (black line) and locally refined RBD/S2D106 variable domains (gray line). The 0.143 cutoff is indicated by a horizontal dashed line. **d**, Local resolution map calculated using cryoSPARC for the whole reconstruction and the locally refined RBD/S2D106 variable domain region. **e,f**, Representative electron micrograph and 2D class averages of SARS-CoV-2 S in complex with the S2H97 Fab embedded in vitreous ice. Scale bar: 400 Å. **g**, Gold-standard Fourier shell correlation curve for the S2H97-bound SARS-CoV-2 S trimer reconstruction. The 0.143 cutoff is indicated by a horizontal dashed line. **h**, Local resolution map calculated using cryoSPARC for the whole reconstruction with two orthogonal orientations.

**Extended Data Fig. 8.**
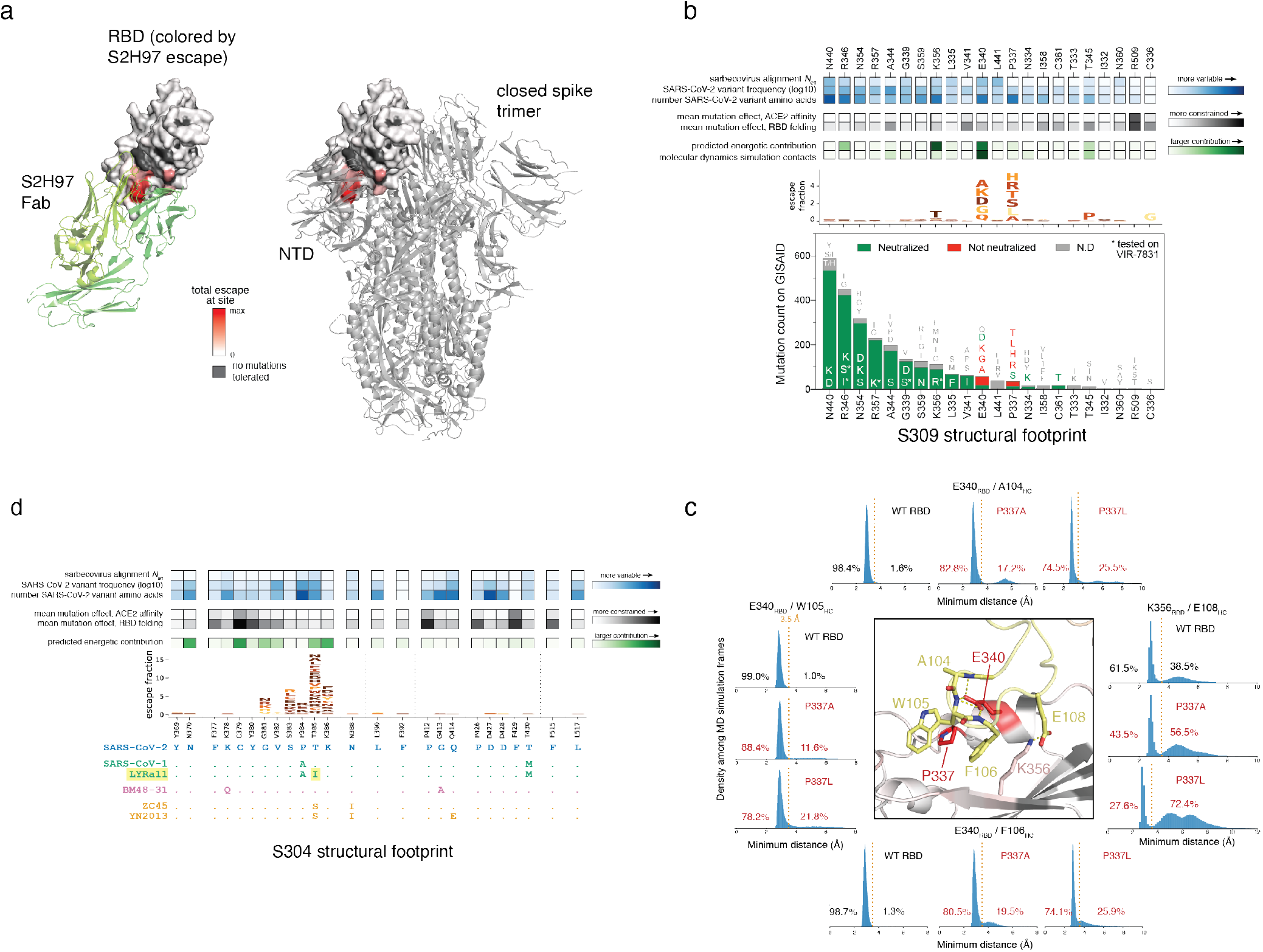
Genetic and structural basis for broad sarbecovirus binding. **a**, Quaternary context of the S2H97 epitope. Left, S2H97-bound RBD, with RBD sites colored by S2H97 escape (scale bar, bottom). Right, RBD in the same angle as left, in the closed spike trimer^40^. The S2H97 epitope is buried with the NTD of the neighboring protomer, requiring extensive RBD opening to enable access to the S2H97 epitope (Fig. 4d). **b**, Integrative genetic and functional features of the structural epitope of S309. Details as in Fig. 3e,f. **c**, Occupancy of key S309/RBD contacts during molecular dynamics simulation of S309 bound to the wildtype (42-μs simulation), P337A (56-μs) and P337L (91-μs) mutated RBD. Details as in Fig. 4g. **d**, Integrative genetic and functional features of the structural epitope of S304. Details as in Fig. 4b.

**Extended Data Table 1.**
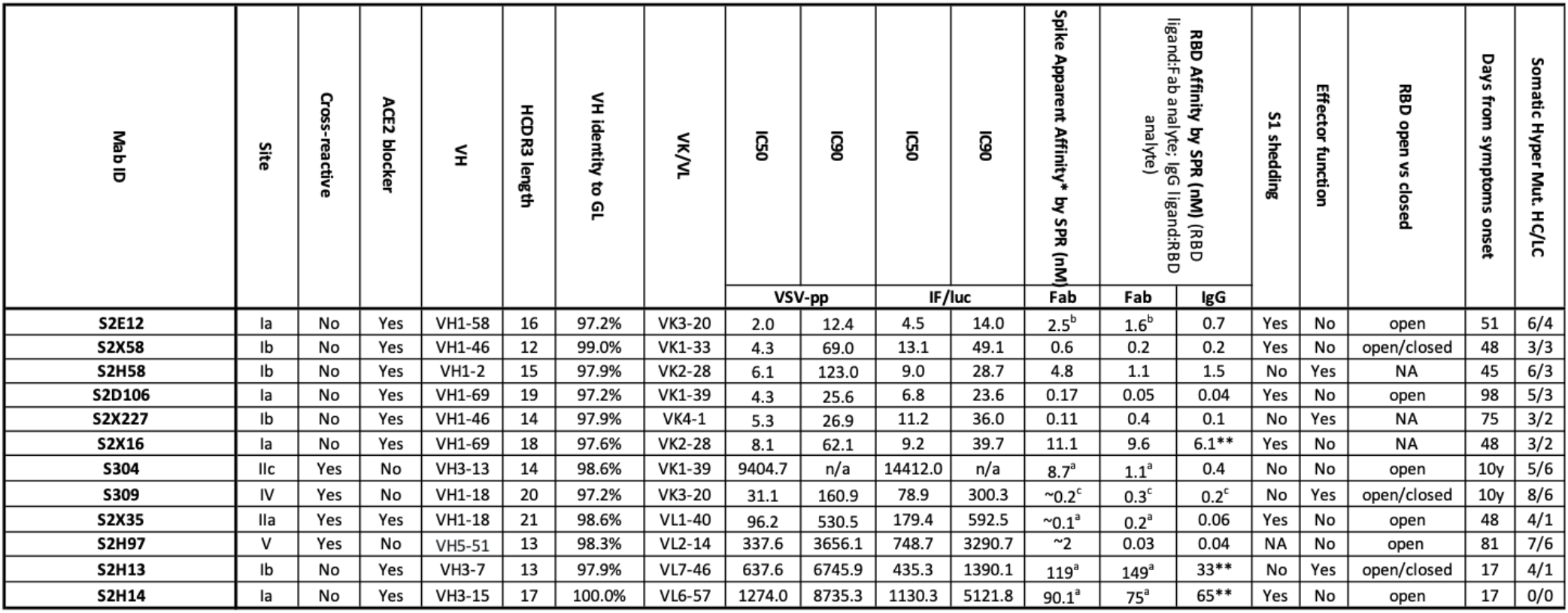
Characteristics of the antibodies described in this study. VH and VL per cent identity refers to V gene segment identity compared to germline (as per the International Immunogenetics Information System (http://www.imgt.org/)). HCDR3 length was determined using IMGT. Binding affinities for previously described antibodies measured in Piccoli et al. (S304, S2X35, S2H13, S2H14)^a^, Tortorici et al. (S2E12)^b^ and Cathcart et al. (S309)^c^. *Spike binding data are “apparent affinity” or KD,app, because RBD conformational dynamics effect the kinetics. S2H97 Fab binding to spike doesn’t fit well to 1:1 binding, presumably because of changing epitope accessibility. **Biphasic kinetics; Fit result is for fast phase

**Extended Data Table 2.**
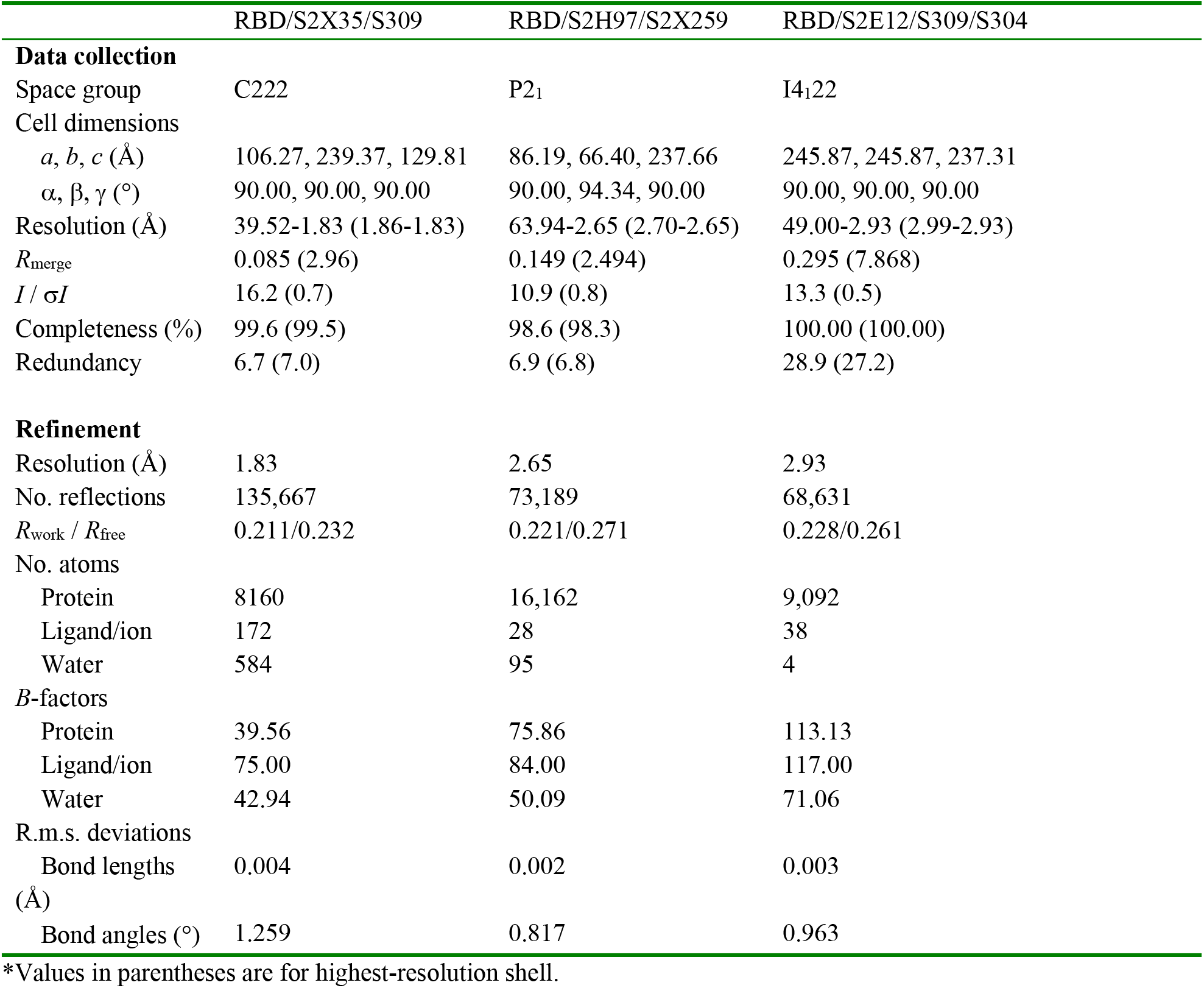
Crystallographic data collection and refinement statistics.

|  | RBD/S2X35/S309 | RBD/S2H97/S2X259 | RBD/S2E12/S309/S304 |
| --- | --- | --- | --- |
| **Data collection** |  |  |  |
| Space group | C222 | P2<sub>1</sub> | I4<sub>1</sub>22 |
| Cell dimensions |  |  |  |
| *a*, *b*, *c* (Å) | 106.27, 239.37, 129.81 | 86.19, 66.40, 237.66 | 245.87, 245.87, 237.31 |
| $\alpha$, $\beta$, $\gamma$ (°) | 90.00, 90.00, 90.00 | 90.00, 94.34, 90.00 | 90.00, 90.00, 90.00 |
| Resolution (Å) | 39.52-1.83 (1.86-1.83) | 63.94-2.65 (2.70-2.65) | 49.00-2.93 (2.99-2.93) |
| *R*<sub>merge</sub> | 0.085 (2.96) | 0.149 (2.494) | 0.295 (7.868) |
| *I* / $\sigma I$ | 16.2 (0.7) | 10.9 (0.8) | 13.3 (0.5) |
| Completeness (%) | 99.6 (99.5) | 98.6 (98.3) | 100.00 (100.00) |
| Redundancy | 6.7 (7.0) | 6.9 (6.8) | 28.9 (27.2) |
| **Refinement** |  |  |  |
| Resolution (Å) | 1.83 | 2.65 | 2.93 |
| No. reflections | 135,667 | 73,189 | 68,631 |
| *R*<sub>work</sub> / *R*<sub>free</sub> | 0.211/0.232 | 0.221/0.271 | 0.228/0.261 |
| No. atoms |  |  |  |
| Protein | 8160 | 16,162 | 9,092 |
| Ligand/ion | 172 | 28 | 38 |
| Water | 584 | 95 | 4 |
| *B*-factors |  |  |  |
| Protein | 39.56 | 75.86 | 113.13 |
| Ligand/ion | 75.00 | 84.00 | 117.00 |
| Water | 42.94 | 50.09 | 71.06 |
| R.m.s. deviations |  |  |  |
| Bond lengths (Å) | 0.004 | 0.002 | 0.003 |
| Bond angles (°) | 1.259 | 0.817 | 0.963 |

\*Values in parentheses are for highest-resolution shell.

